# Form, function, and divergence of a generic fin shape in small cetaceans

**DOI:** 10.1101/2020.07.21.214734

**Authors:** Vadim Pavlov, Cecile Vincent, Bjarni Mikkelsen, Justine Lebeau, Vincent Ridoux, Ursula Siebert

## Abstract

Tail flukes as well as the dorsal fin are the apomorphic traits of cetaceans appeared during evolutionary process of adaptation to the aquatic life. Both appendages present a generic wing-like shape associated with lift generation and low drag. Variability of the form of appendages was studied in seven species of cetaceans having different body size, external morphology, and specialization. Hydrodynamic performance of the fin cross-sections was examined with the CFD software and compared with similar engineered airfoils. Affinity of hydrodynamic design of both appendages was found in a wing-like planform and cross-sectional design optimized for lift generation. Distinctions in the planform and cross-sections were found related with the fin specialization in thrust production or swimming stability control. Cross-sectional design of the dorsal fin was found to be optimized for the narrow range of small angles of attack. Cross-sections of tail flukes were found to be more stable for higher angles of attack and had gradual stall characteristics that is associated with their propulsive efficiency as oscillating foils. The results obtained are the evidence of divergent evolutionary pathways of a generic wing-like shape of the fins of cetaceans under specific demands of thrust production and swimming stability control.

## Introduction

Question of the role of dolphin appendages as lift-generating surfaces is related to the evolutionary process of adaptation of marine mammals to the life in “a moving fluid” [1]. In this context, the dorsal fin and tail flukes of cetaceans are of particular interest, as there is no evidence of their analogs in terrestrial ancestors [2–4] and their appearance in cetaceans is presumably associated with transition from drag-based to lift-based locomotion in an aquatic environment [5, 6]. As *de novo* dermal structure [7], the dorsal fin and tail flukes can be described with a limited set of morphological traits where the relation between the traits and wing performance can be unambiguously interpreted. This unambiguous interpretation provides for insight into the evolutionary pathways to divergence of a generic shape driven by the different role in swimming stability control and thrust production.

Both appendages represent an underwater wing, where the fin span (*S*) and fin planform area (*A*) correlate with the body length (*BL*) of cetaceans [4, 8, 9]. Morphology and hydrodynamic design of appendages are well adapted for high lift and low drag [1]. Larger wings generate more lift than wings of smaller area or span and are associated with increased thrust production and stability control of large species when swimming. The relationships between *S*, *A*, and *BL* is different in life history stages that affect the swimming performance of cetaceans [8, 9].

The planform of tail flukes most often presents a falcate, swept-back tapered outline, with moderate or high aspect ratio *AR* = *S*^2^/*A* [4, 8, 9]. The dorsal fins of the variety of species of cetaceans normally have lower *AR* and more variable planforms with positive, neutral, and negative sweep of the trailing edge that appears as falcate, rounded, and triangle shape of the fins [10]. Cross-sectional design of both dorsal fins and flukes displays a symmetrical streamlined outline with rounded leading edge [11–16]. This shape is comparable with the engineered airfoils and hydrofoils [11, 15, 16, 17].

The combination of moderate aspect ratio, sweep, cross-sectional design, and flexibility of the fins characterizes efficient underwater wing [18–21]. Meanwhile, there is a fundamental difference in the operational mode of the dorsal fin as a fixed wing and the tail flukes acting as a pair of oscillating wings [1, 11, 16, 20]. Apart from the fixed wing, an oscillating wing is involved in specific mechanisms of lift and drag generation dealing with leading edge vortex and wake capture [22]. The advantage of an oscillating mode is low drag, increased lift, delayed stall and wider range of the angles of attack [1, 23, 24].

Here, we investigated morphological variability of the dorsal fins and tail flukes of small cetaceans and connected it to their role in swimming stabilization control and thrust production. We carried out interdisciplinary quantitative approach to reveal the affinity and distinctions of fins related with their specialization. Using the combination of morphometric and hydrodynamic analysis, we present the evidence of divergent evolutionary pathways of a generic wing-like shape of the fin of cetaceans.

## Results

### Size, shape, and cross-sectional geometry of the dorsal fins

Both size and shape of the dorsal fin varied significantly among the species studied. Fin span varied from *S* = 10 ± 1.5 (means ± SD) cm in harbor porpoise to *S* = 30.4 ± 7.1 (means ± SD) cm in pilot whale. Span *S* and area *A* of the fin increased with increasing length of the body (Fig 1). Distinctions of the fin shape were revealed in relative parameters *AR* and *CI* (Table 1) as well as in sweep of the trailing edge of the fin. Fin shape varied from triangle one with low *AR* and positive sweep of the trailing edge in harbor porpoise to the falcate shape with moderate or high *AR* and negative sweep of the trailing edge in common dolphin, bottlenose dolphin, Atlantic white-sided dolphin, white-beaked dolphin, and Minke whale. Within that group the relative parameters *AR*, *Cl*, and *Λ* varied moderately (Table 1). Apart from these species the dorsal fins of harbor porpoise and pilot whale had obvious distinctions in absolute and relative parameters of the fin shape.

**Fig 1.**
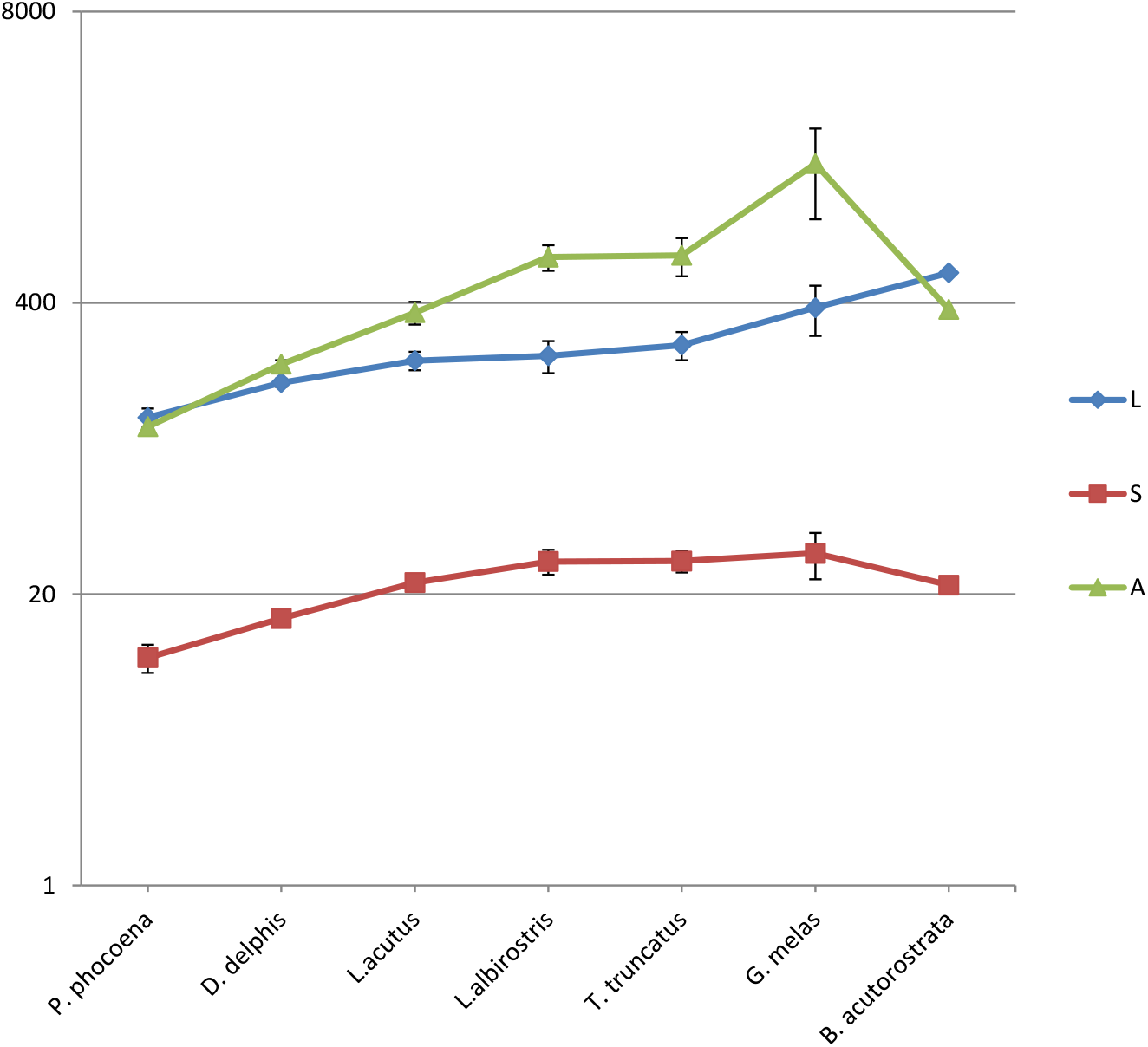
Species-specific differences in the length of the body *L* cm, span of the dorsal fin *S* cm, and area of the dorsal fin *A* cm^2^, means ± SD.

**Table 1.**
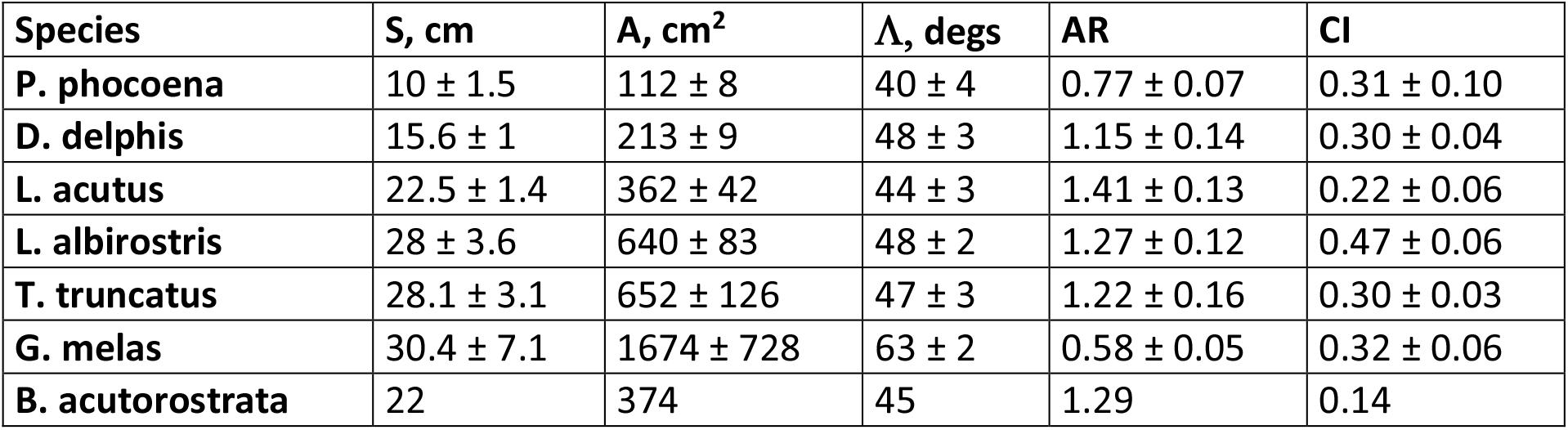
Absolute and relative parameters of the dorsal fins, means ± SD. Species are ranged by the body size.

Cross-sections of dorsal fins were close to the conventional symmetrical airfoils by Eppler, Selig, and NACA [25]. Cross-sections at the base of dorsal fins were comparable with laminarized profiles E297 and E836 with the thickness ratio increased to 15%, while the cross-sections at the fin tip had a similarity with the E477, S1048, and NACA 0015 airfoils. A smooth transition of shape of cross-section taken from the bottom and tip of the fin was observed. At noticeable similarity to the airfoils geometry all fin cross-sections had distinctively thickened trailing edge.

Absolute parameters of fin cross-sections *CL, MT*, and *PMT* related with the fin size varied in wide range with the extreme values for harbor porpoise and Minke whale (Figs S1 – S3). Meanwhile the span-wise variation of these parameters appeared similar in all species studied. The geometry of cross-sections at the base of the fin was more variable compared with cross-sections located at the tip of the fin.

Apart from absolute parameters of cross-sections, the relative ones appeared to be more consistent (Figs S4 – S6). From fin base to the fin tip the *MT*, %*CL* decreased in pilot whale and white-beaked dolphin, varied slightly in bottlenose dolphin, Atlantic white-sided dolphin and Minke whale, and increased in common dolphin and harbor porpoise. In same direction *PMT*, %*CL* varied slightly in harbor porpoise, white-beaked dolphin and Minke whale, and increased in common dolphin, Atlantic white-sided dolphin, bottlenose dolphin, and pilot whale.

Span-wise distribution of *r*, %*CL* was revealed similar in all species studied. In general, this parameter increased from fin base to the two thirds of the fin span in all species, then varied slightly to the fin tip. No species-specific distinctions were observed for this parameter, except of the pilot whale, where average values of *r*, %*CL* were significantly higher compared with other species.

### Size, shape, and cross-sectional geometry of the flukes

With the variable size, *S* = 30 ± 3 (means ± SD) cm in harbor porpoise and *S* = 165 cm in Minke whale, the shape of the tail flukes appeared to be less variable compared with shape of the dorsal fins (Table 2). All species had positive sweep of leading and trailing edge where *Λ* of the leading edge varied from 27 degrees in Minke whale to 35 ± 1 (means ± SD) degrees in bottlenose dolphin. Distinctions were revealed between the pilot whale and Minke whale group having trapezoidal shape of the tail flukes with high *AR*, and Atlantic white sided dolphin, bottlenose dolphin and harbor porpoise group having swept back tips of the flukes and lower *AR* (Table 2). Common dolphin and white-beaked dolphin had moderate sweep back of the tips of the flukes. Both *S* and *A* of the flukes had positive correlation with the length of the body (Fig 2).

**Fig 2.**
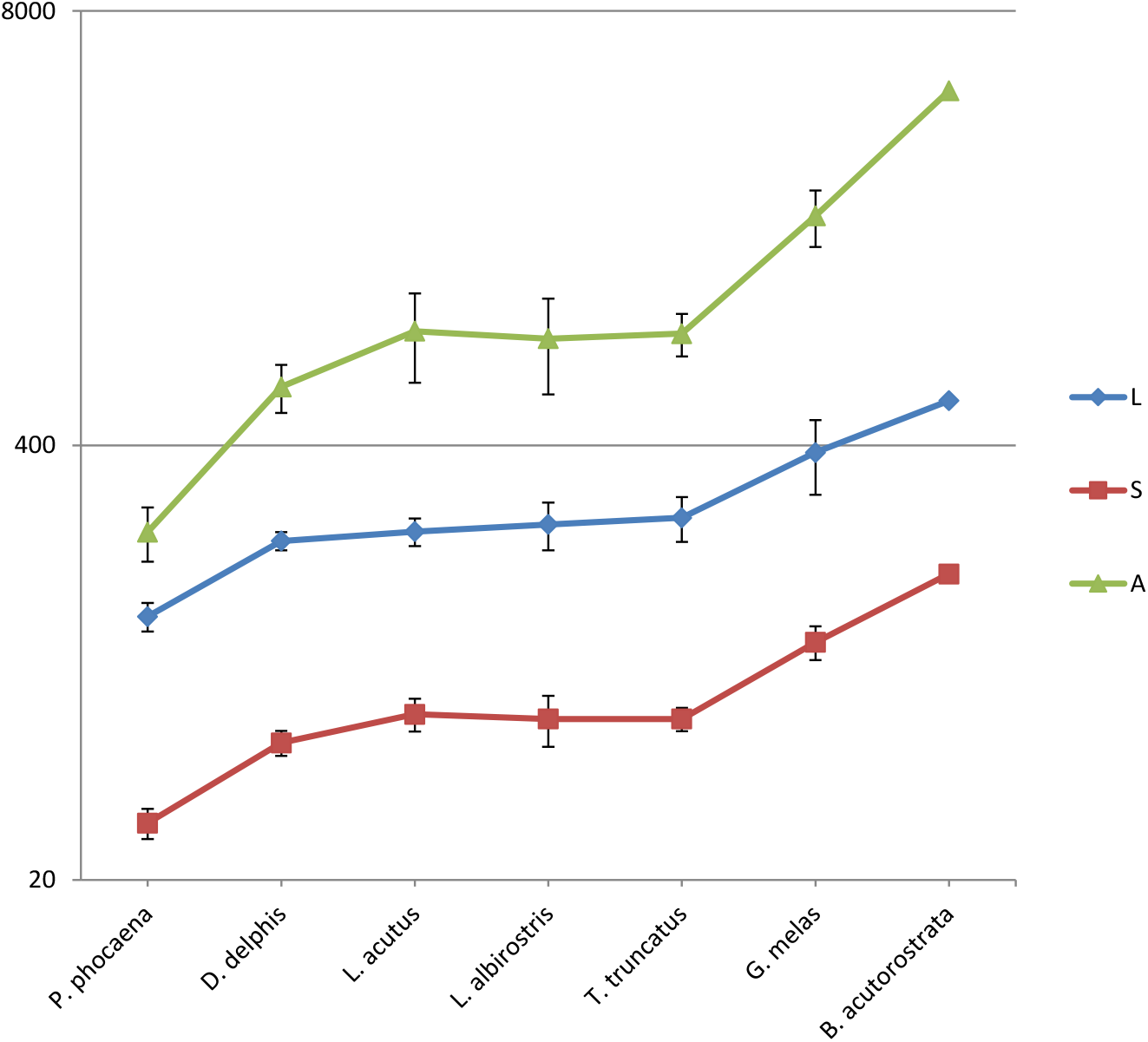
Species-specific differences in the length of the body *L* cm, span of the fluke *S* cm, and area of the fluke *A* cm^2^, means ± SD.

**Table 2.**
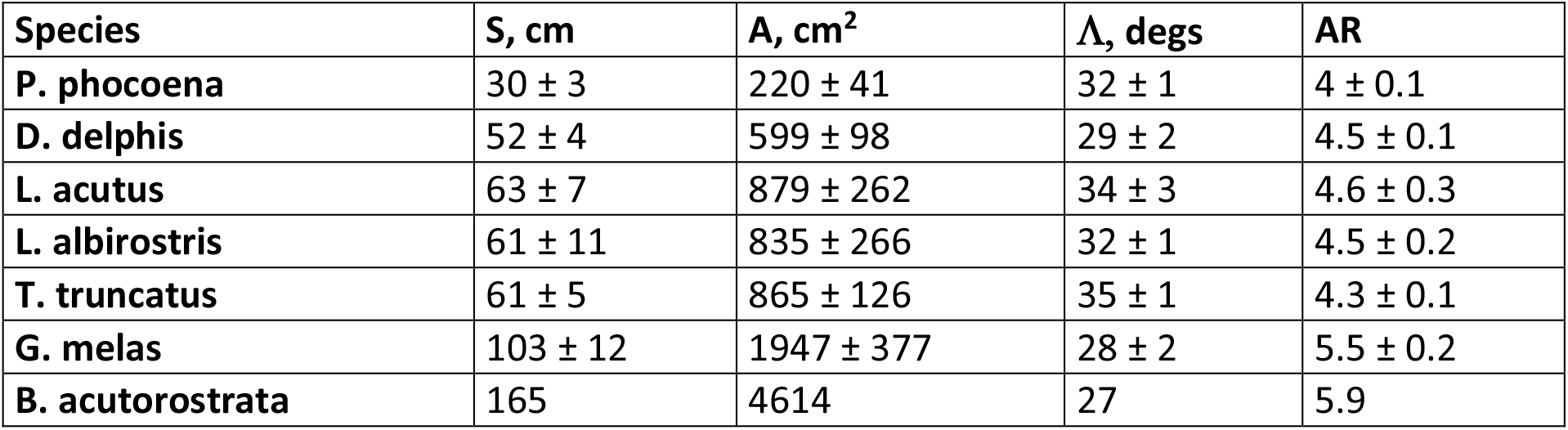
Absolute and relative parameters of the flukes, means ± SD. Species are ranged by the body size.

Fluke’s cross-sections had a resemblance with the conventional symmetrical airfoils by Eppler, Selig, and NACA [25]. Absolute parameters *CL, MT*, and *PMT* varied insignificantly in common dolphin and bottlenose dolphin (Figs S7 – S9). Minimal and maximal values referred to the harbor porpoise and bottlenose dolphin respectively. All species had similar span-wise distribution of *CL, MT*, and *PMT* decreasing from fin base to the fin tip. Except of the harbor porpoise, cross-sections located at the base of the flukes possessed most variable parameters *CL, MT*, and *PMT*.

Relative parameters of tail fluke cross-sections varied less than the absolute ones (Figs S10 – S12). Similar pattern of *MT*, %*CL*, *PMT*, %*CL*, and *r*, %*CL* distribution in span-wise direction was found in all species studied. Relative thickness of the cross-sections *MT*, %*CL* decreased abruptly from the bottom of fin to the 17% of the fluke’s span and then changed slightly to the fluke tip in common dolphin and bottlenose dolphin. In harbor porpoise this parameter decreased more obviously to the fluke tip.

Position of relative thickness *PMT*, %CL in fluke’s cross-sections increased from the bottom to the tip of the flukes. Average values of *r*, %*CL* on the cross-sections at the bottom of the flukes differed by the order from these ones on other cross-sections. In general, *r*, %*CL* decreased from the bottom of the flukes to the mid-span and then gradually increased to the tip of the flukes.

Peculiarities in cross-sectional geometry of the dorsal fin and tail fluke were examined with the Principal Component Analysis (Fig 3). Distinctive separation between the dorsal fin and fluke’s cross-section dimensionless parameters was found, where the main difference between the dorsal fin and fluke’s cross-sectional geometry was associated with higher *LER*%*CL* in fluke’s cross-sections and higher *PMT*%*CL* in the dorsal fin cross-sections.

**Fig 3.**
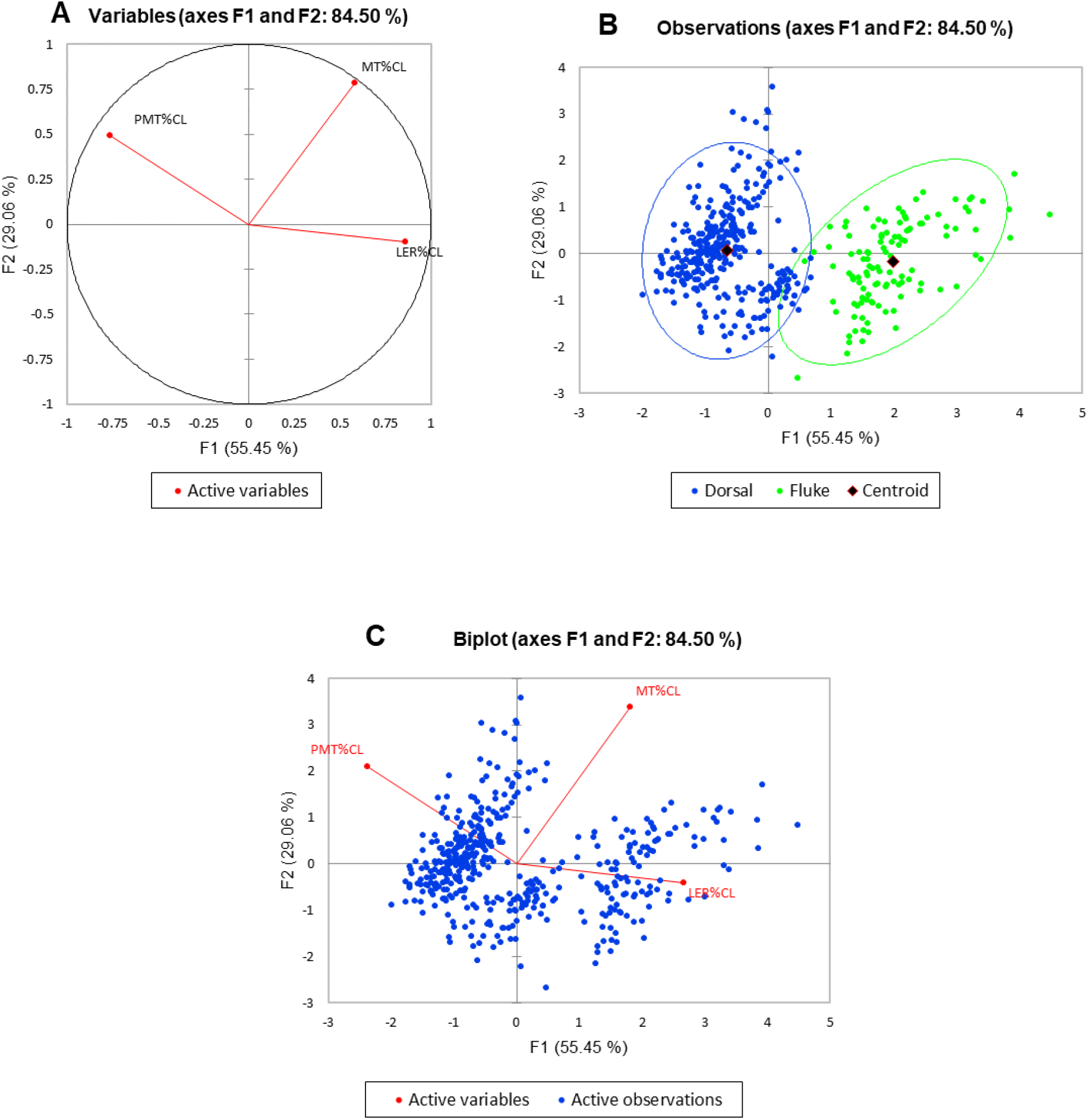
Principal Component Analysis of the dimensionless parameters of the dorsal fin and tail fluke cross-sections of all species. A – Negative relationship between *LER*%*CL* and *PMT*%*CL*, while the *MT*%*CL* appears to be unrelated with the *LER*%*CL* and *PMT*%*CL*. B – Separation between the dorsal fin and fluke’s cross-section dimensionless parameters. C – The main difference between the dorsal fin and fluke’s cross-sectional geometry is associated with higher *LER*%*CL* in fluke’s cross-sections and higher *PMT*%*CL* in the dorsal fin cross-sections.

### Hydrodynamic performance of the cross-sections of dorsal fin

At routine speed of swimming 2 m/sec [10] the drag coefficient *Cd* of the fin cross-sections increased from the fin bottom to the fin tip (S13 Fig). At burst speed of swimming 8 m/sec the *Cd* slightly decreased while a span-wise pattern of *Cd* distribution remained the same (S14 Fig).

Average values of *Cd* of the cross-sections located at the bottom of the fin were comparable with the *Cd* of E297 and E836 airfoils calculated under the same conditions, see Table 3. Pressure distribution had sharp negative pressure gradient at the leading edge with minimal *Cp* values at 15-20 % of the chord length. Both in cross-sections and airfoils the region from 20 to 60 % of chord length was characterized by zero or slightly positive pressure gradient.

**Table 3.** Comparison of the hydrodynamic performance of the dorsal fin cross-sections with the appropriate airfoils.

| Species | Section # | Re number | Cd 2 m/sec | Re number | Cd 8 m/sec | min Cp |
| --- | --- | --- | --- | --- | --- | --- |
| P. phocoena | 1 | 3,6E+05 | 0,0100 | 1,5E+06 | 0,0073 | -0,383 |
| D. delphis | 1 | 4,4E+05 | 0,0083 | 1,8E+06 | 0,0065 | -0,398 |
| T. truncatus | 1 | 5,3E+05 | 0,0101 | 2,7E+06 | 0,0077 | -0,456 |
| L. acutus | 1 | 5,3E+05 | 0,0089 | 2,1E+06 | 0,0070 | -0,414 |
| G. melas | 1 | 9,2E+05 | 0,0110 | 3,7E+06 | 0,0081 | -0,514 |
| L. albirostris | 1 | 5,5E+05 | 0,0101 | 2,9E+06 | 0,0073 | -0,398 |
| M |  | **5,6E+05** | **0,0097** | **2,4E+06** | **0,0073** | **-0,427** |
| SD |  | 1,9E+05 | 0,0010 | 8,0E+05 | 0,0006 | 0,049 |
| **airfoils** |  |  |  |  |  |  |
| E297* |  | **5,6E+05** | **0,0081** | **2,4E+06** | **0,0062** | **-0,416** |
| E836* |  | **5,6E+05** | **0,0072** | **2,4E+06** | **0,0055** | **-0,405** |
| **Species** |  |  |  |  |  |  |
| P. phocoena | 6 | 1,4E+05 | 0,0135 | 5,7E+05 | 0,0090 | -0,515 |
| D. delphis | 6 | 1,7E+05 | 0,0136 | 6,6E+05 | 0,0103 | -0,572 |
| T. truncatus | 6 | 2,1E+05 | 0,0124 | 7,7E+05 | 0,0092 | -0,488 |
| L. acutus | 6 | 1,8E+05 | 0,0132 | 7,1E+05 | 0,0094 | -0,507 |
| G. melas | 6 | 3,2E+05 | 0,0106 | 1,3E+06 | 0,0085 | -0,443 |
| L. albirostris | 6 | 1,8E+05 | 0,0111 | 7,1E+05 | 0,0092 | -0,524 |
| M |  | **2,0E+05** | **0,0124** | **7,8E+05** | **0,0093** | **-0,508** |
| SD |  | 6,3E+04 | 0,0013 | 2,5E+05 | 0,0006 | 0,042 |
| **airfoils** |  |  |  |  |  |  |
| E477* |  | **2,0E+05** | **0,0113** | **7,8E+05** | **0,0084** | **-0,486** |
| S1048* |  | **2,0E+05** | **0,0121** | **7,8E+05** | **0,0087** | **-0,532** |

Average values of *Cd* of cross-sections located at the tip of the fin were comparable with the *Cd* of E477 and S1048 airfoils calculated under the same conditions (Table 3). Peculiarity of pressure distribution there was an absence of zero or slightly positive pressure gradient. Change of the sign of gradient occurred at 20-25% of chord length.

Laminar region on the cross-sections ranged from 20 to 28 % of the chord length, while the length of transition zone varied from 5 to 30 % of the chord length. Relative laminar flow portion of the boundary layer, % *CL* had a general trend to decrease from fin bottom to the fin tip. This regularity was found common for all species studied except of harbor porpoise (S15 Fig). Reduction of the laminar flow portion related with increasing speed of swimming to 8 m/sec was observed for all species (S16 Fig).

### Hydrodynamic performance of the cross-sections of tail flukes

At routine speed of swimming 2 m/sec the *Cd* of the fluke cross-sections decreased from fluke’s bottom to 17-34% of the fluke’s span and then gradually increased to the fluke tip (S17 Fig). At burst speed of swimming 8 m/sec the *Cd* of the fluke’s cross-sections decreased while a span-wise pattern of *Cd* distribution remained the same (S18 Fig).

Average *Cd* of cross-sections was comparable with the *Cd* of SD8020 and S8035 airfoils calculated under the same conditions (Table 4). Pressure distribution along the cross-sections located at the base of fluke was close to mentioned above SD8020 and S8035 airfoils. Peculiarity of these profiles was the absence of zero or slightly positive pressure gradient. Due to higher curvature of cross-sections the length of transition zone varied in narrow range from 5 to 8% of *CL*. Specific shape of cross-sections at the base of fluke had lowest minimal values of pressure coefficient *Cp* −1.073 that exceeded twice the value 0.427 of the same parameter of cross-sections located at the base of the dorsal fin.

**Table 4.** Comparison of the hydrodynamic performance of the tail fluke cross-sections with the appropriate airfoils.

| Species | Section # | Re number | $C_d$ 2 m/sec | Re number | $C_d$ 8 m/sec | min $C_p$ |
| --- | --- | --- | --- | --- | --- | --- |
| P. phocoena | 1 | 4,3E+05 | 0,0109 | 1,7E+06 | 0,0097 | -1,053 |
| D. delphis | 1 | 6,0E+05 | 0,0115 | 2,4E+06 | 0,0085 | -1,120 |
| T. truncatus | 1 | 7,9E+05 | 0,0122 | 3,2E+06 | 0,0087 | -1,047 |
| M |  | **6,1E+05** | **0,0115** | **2,4E+06** | **0,0090** | **-1,073** |
| SD |  | 1,8E+05 | 0,0007 | 7,3E+05 | 0,0006 | 0,041 |
| **airfoils** |  |  |  |  |  |  |
| S8035* |  | **6,1E+05** | **0,0117** | **2,4E+06** | **0,0084** | **-1,021** |
| SD8020* |  | **6,1E+05** | **0,0115** | **2,4E+06** | **0,0087** | **-1,019** |
| **Species** |  |  |  |  |  |  |
| P. phocoena | 6 | 1,4E+05 | 0,0128 | 5,6E+05 | 0,0094 | -0,515 |
| D. delphis | 6 | 1,7E+05 | 0,0116 | 6,6E+05 | 0,0094 | -0,576 |
| T. truncatus | 6 | 2,0E+05 | 0,0124 | 7,8E+05 | 0,0090 | -0,473 |
| M |  | **1,7E+05** | **0,0123** | **6,7E+05** | **0,0093** | **-0,521** |
| SD |  | 3,0E+04 | 0,0006 | 1,1E+05 | 0,0002 | 0,052 |
| **airfoils** |  |  |  |  |  |  |
| E477* |  | **2,0E+05** | **0,0113** | **7,8E+05** | **0,0084** | **-0,486** |
| S1048* |  | **2,0E+05** | **0,0121** | **7,8E+05** | **0,0087** | **-0,532** |

\*Modified profile with thickness ratio increased to 15%.

The shape of cross-sections located at the tip of the flukes was quite similar to the cross-sections of the dorsal fin made at the same locations (Tables 3, 4). As a consequence, average *Cd* as well as pressure distribution of the cross-sections at that location differed slightly (Table 4).

Laminar region on the cross-sections ranged from 18 to 32% of the chord length while the length of transition zone varied from 5 to 20% of chord length. Relative laminar flow portion of the boundary layer, % *CL* varied slightly in span-wise direction (S19 Fig). As in the dorsal fin, reduction of the laminar flow portion associated with increasing speed of swimming up to 8 m/sec was observed for all species studied. In common dolphin, the laminar flow portion gradually decreased from fluke base to the tip, while in harbor porpoise and bottlenose dolphin this parameter increased from fluke base to 17% of the fluke’s span and then changed insignificantly (S20 Fig).

### Comparison of hydrodynamic performance of the dorsal fin and flukes

Although both dorsal fin and fluke had similar cross-sectional design, they had specific distinctions influenced their hydrodynamic performance. In general, the cross-sections of dorsal fin were thinner, had smaller *r*, %*CL*, and the *PMT*, %*CL* decreased from the fin base to the fin tip apart from increased *PMT*, %*CL* in the same direction in tail flukes.

Cross-sections of both appendages exhibited the lowest drag over a narrow range of angle of attack called the “drag bucket” for simulated cruising speed of swimming 2 m/sec (S21 and S23 Figs). The term “drag bucket” is used to describe the shape of a drag curve showing Cd against α where the drag curve shows an extended flat bottom of the curve i.e. bucket shaped [17]. The shape and position of drag bucket was solely determined by the cross-sectional geometry. Generally, the width of drag bucket decreased from the bottom to mid-span of the fin and then increased to the fin tip, according to the span-wise distribution of *r*, %*CL, MT*, %*CL* and the *PMT*, %*CL*.

Main difference as in geometry as in hydrodynamic performance of cross-sections was related to the fin-body junction. Cross-sections of dorsal fin of the Atlantic white-beaked dolphin located at this region possessed the highest L/D ratio for the narrow range of α = 05° and least stall angle 8° (S21 and S22 Figs). These cross-sections had more abrupt stall characteristics compared with the mid-span and fin tip location. Cross-sections of the fluke of common dolphin taken at the same location had the best L/D ratio among fluke cross-sections and stall angle 14° (S23 and S24 Figs).

Span-wise lift distribution *C*Cl_max_* on cross-sections of both appendages presented some approximation of the optimal elliptical one with decreasing L from the bottom of the fin to the fin tip (S25 Fig). Absolute values of this parameter on the dorsal fin cross-sections were approximately two times higher compared with the fluke cross-sections.

## Discussion

Dorsal fins and tail flukes of small cetaceans present an example of a natural underwater wing well adapted for lift production at low drag. This example is interesting from the phylogenetic evolution point of view as both fins appeared as highly specialized organs in the absence of evidence of their analogs in terrestrial ancestors of cetaceans. Distinctive feature of these apomorphic structures is an unambiguous relation between their wing-like shape and function in lift production. Morphological distinctions of dorsal fin and tail flukes were found to be closely associated with their different role as a vertical keel for stability control (dorsal fin) and oscillating foil generating lift-derived thrust (flukes).

### Hydrodynamics of fins

Hydrodynamic performance of a wing can be estimated from the ratio of lift-to-drag generated when moving in a fluid [26]. The wings, keels, and control surfaces in sea and air vehicles are designed for specific purposes that anticipate the optimal wing performance within certain range of operational conditions [27, 28, 29]. Optimization of a wing for the expected range of speeds and angles of attack can be done by the altering the basic wing and foil parameters including span, area, sweep, twist, taper, *r*, *MT, PMT*, and a camber. Analysis of the natural wings design could help better understanding their specialization and the range of operational conditions for which they are optimized.

Both dorsal fin and tail flukes have symmetric cross-sectional design that generates no lift at α = 0. When the fin is canted to the flow at certain α, the pressure differences on both sides of the fin generate lift and lift-induced drag associated with the span-wise crossflows that go around the fin tip [17]. The magnitude of the induced drag is determined by the span-wise lift distribution and the resulting distribution of vortices [28]. In general, the fins have the swept-back tapered planform and span-wise variability of cross-sectional geometry, which is known as aerodynamics twist in wing engineering practice [17, 31]. Both these features may influence lift distribution so that more lift can be generated at the wing root and less towards the wingtip (S25 Fig) that causes a reduction in the strength of the wingtip vortices and reduction of lift-induced drag [28]. The bending of fin both in span-wise and chord-wise direction also limits the lift and therefore lift-induced drag [31]. Both kinds of appendages had smooth filleted joint with the dolphin’ body. That feature of dolphin anatomy may facilitate decreasing the interference drag appearing when surfaces at angles to each other simulate turbulence in the region of the joint as it can be observed in the intersection of the fuselage and wing in aviation [32].

The morphology and biomechanics of flukes have distinctive features associated with drag reduction and generation of lift-derived thrust. The high aspect ratio platform of the flukes is advantageous in producing less induced drag compared with low aspect ratio dorsal fin [28]. The mode of flukes bending varies from uniform bend of the entire fluke’s blades to the characteristic shape of blended winglets of the airliners’ wings [33, 34]. The latter one is related with reduction of induced drag by decreasing the vortex formation on the wingtip area [34, 35].

Clear distinctions in root cross-section geometry are likely to be related with the principal difference between static and oscillating hydrofoils. The static wing performance is limited by the angles of attack causing stall condition, i.e. dramatic decrease of the lift and increase drag. Oscillating wings, on the contrary, can operate within wider range of the angles of attack without loss of efficiency because of dynamic stall caused by unsteady effects [22, 23, 36]. In the species studied, the dorsal fin had extended and thin root cross-section with small leading edge radius. This design is widespread in aircraft vertical stabilizer, e.g. Boeing 737, as well as in yacht keel design and associated with least interference drag, best performance within narrow range of α, but sudden stall characteristics outside the narrow lift-coefficient range [28, 29]. Apart from that, thick root cross-section of the flukes with large leading edge radius and shifted forward *PMT* permits greater angle of attack variations without producing excessive minimum pressure peaks at the leading edge that can result in flow separation and cavitation [17]. This difference in optimization of the general foil shape to the static and oscillating wing function is expressed in different lift, drag, and stall characteristics of root cross-section of the dorsal fin and tail flukes as shown in drag polar diagrams (S Figs 21-24).

### Optimization of a generic shape to the specific function

Transition from terrestrial to the aquatic environment in cetacean ancestors was accompanied by the numerous morphological and physiological adaptations including changes in their musculoskeletal system [37, 38, 39]. Modern cetaceans have pectoral fins homological to the forelimbs of terrestrial mammals and rudimentary hind limbs. Transition to the aquatic locomotion which is biologically propelled motion through a liquid medium has led to appearance of tail flukes generating lift-derived thrust [40]. Dorsal fin, if present, appeared as vertical stabilizer preventing rolling and side-slip. Apart from pectoral fins, the tail flukes and dorsal fin are the apomorphic traits in cetaceans along with skin blubber, isolated ear bone, increased vertebrae, and other adaptations [39]. These appendages have no homology in cetacean terrestrial ancestors and are interesting as *de novo* wing-like structures [7, 4] specialized for lift production.

Comparison of tail flukes and dorsal fin in cetaceans presents an interesting example of the pathways of evolutionary novelty development within certain taxonomic group. The morphological traits of a generic wing-like shape present conventional wing parameters that facilitates their interpretation in terms of lift and drag. Variability of the morphological traits is restricted by the wing design as well as by structural limitation to the strength and stiffness of the fins [14, 41]. This limited variability, though, may be different in some specific directions related with a wing function. The difference in variation of *r* and *PMT* of the fin cross-section is associated with the involvement of tail flukes and dorsal fin in the opposite scenarios of thrust production and swimming stability control.

Morphological constraints of a generic wing-like shape are also appearing to be strong or weaken depending on the different demand in wing performance. The tail flukes being the only organ of locomotion in cetaceans show consistency in wing planform including inverse relationship between *Λ* and *AR* across the representatives of both Odontoceti and Mysticeti [4]. As the lift produced by wing is proportional to the wing area the relation between *S* and *A* of the flukes and body length of a cetacean is expectable. It can be observed as in different taxa as in different life history stages [8, 9, 4].

Apart from the flukes, the swimming stabilization control is not solely the function of dorsal fin as it is shared between other appendages as well as the tail peduncle [42]. The shape of the dorsal fin is much more variable and may be related with age and sex of cetaceans [8, 10, 43, 44]. The relation between *S* and *A* of the fin and body length is observed in this article for the species known as good swimmers but this is not common for all cetaceans. Both *S* and *A* of the dorsal fin vary dramatically from high *AR* fin in killer whale to low *AR* fin in harbor porpoise, dorsal ridge in river dolphins, and absence of fin in finless species such as finless porpoise *Neophocaena phocaenoides* or Northern right whale dolphin *Lissodelphis borealis*. A spread falcate shape of dorsal fin can be observed across the different families of cetaceans but the contribution of fin into stability control seems to be different in a 2 m dolphin and a baleen whale with much larger size of the body (Fig 1).

Comparison of the tail flukes and dorsal fin variability shows the evolutionary pathways of a generic wing-like shape of the fin as an apomorphic trait of cetaceans. High demand in performance led to high specialization and consistent morphological design of propulsive appendages. Absolute parameters *S* and *A* of the tail flukes are related with the increasing body length of Odontocetes as both the span and area determine the mass of water that is affected for thrust generation [4]. Apart from that, the lower demand in the dorsal fin performance resulted in much more variable design and led to reduction or even elimination of the fin morphology and function.

The results obtained can stimulate the real-world applications of three-dimensional oscillating foils where the effects of three-dimensionality, free stream turbulence and high Reynolds numbers can be considered. The data obtained may serve as a baseline in construction of propulsors and control surfaces for AUV’s operating within the similar range of Reynolds numbers 10^5^ – 10^7^. It could also facilitate optimization of the external design of the fin-mounted tags for small cetacean telemetry.

## Materials and Methods

### Sampling

Dorsal fins and tail flukes in good condition were taken from dead stranded and by-caught animals in the Bay of Biscay, Black Sea, North Sea, and Norwegian Sea. Measurements of the body length and dorsal fin were taken from 3 harbor porpoises *Phocoena phocoena*, 16 common dolphins *Delphinus delphis*, 10 bottlenose dolphins *Tursiops truncatus*, 11 Atlantic white-sided dolphins *Lagenorhynchus acutus*, 3 Atlantic white-beaked dolphins *Lagenorhynchus albirostris* and 13 long-finned pilot whales *Globicephala melas* (Fig 4). Measurements of the flukes were taken from 3 *P. phocoena*, 4 *D. delphis*, 3 *T. Truncatus*, 3 *L. acutus*, 3 *L. albirostris*, and 3 *G. melas*. Measurements were made on one fluke, left or right, depending on the condition. Additionally, the measurements of the dorsal fin and flukes were taken from one specimen of the Minke whale *Balaenoptera acutorostrata*. Fin span was measured from notch point to the fin tip in tail flukes and from the root chord to the top in dorsal fin. The position of root chord was assumed a line parallel to the long axis of the body passing through the point of the maximal curvature on the leading edge at the base of the fin. Fins were cut off from the body and six cross-sections parallel to the long axis of the body were made in equal interval. Photographs of the intact fins and cross-sections were taken with the ruler for scale.

**Fig 4.**
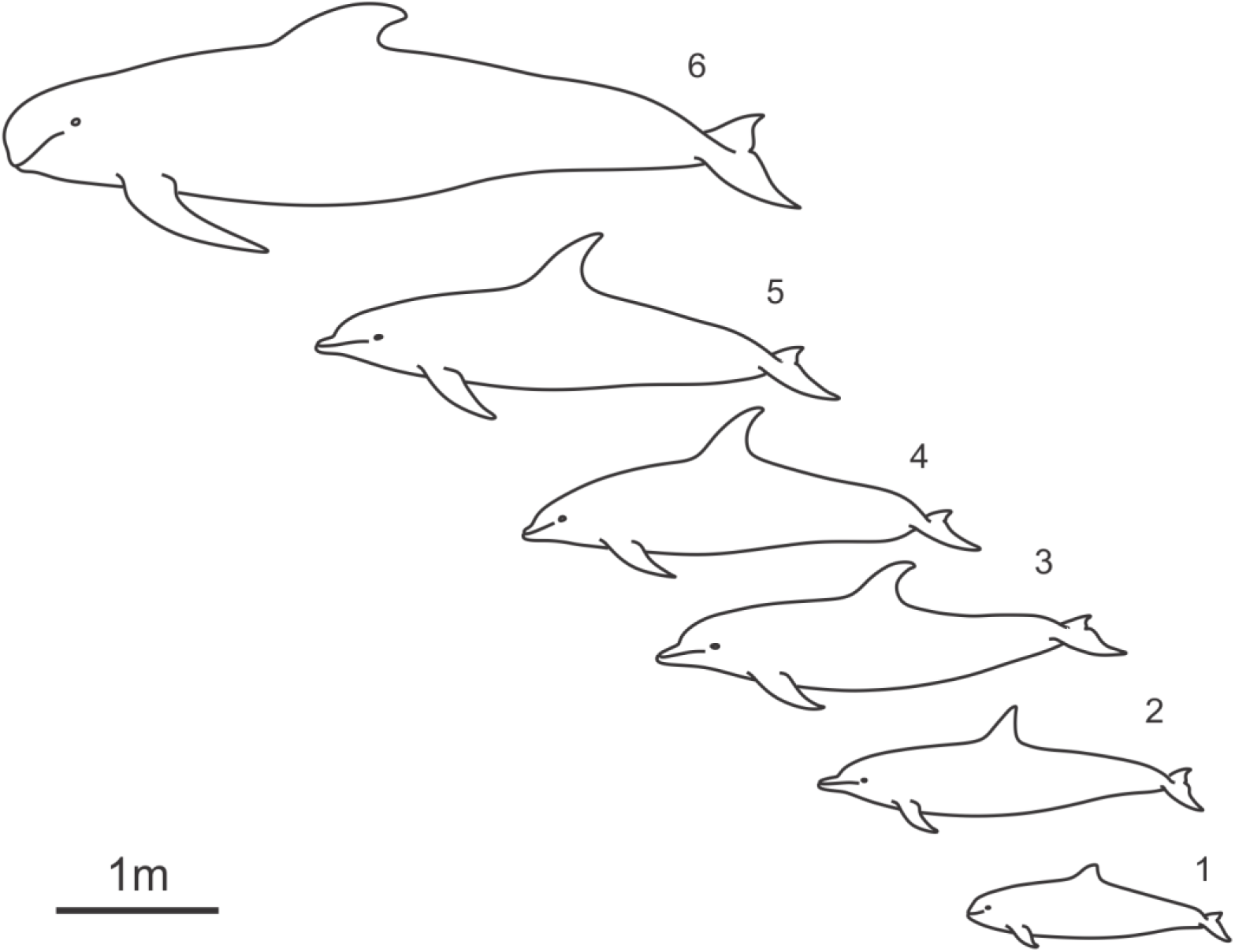
Small cetacean species selected for this study. 1 – *Harbor porpoise*, 2 – *Delphinus delphis*, 3 – *Tursiops truncatus*, 4 – *Lagenorhynchus acutus*, 5 – *Lagenorhynchus albirostris*, 6 – *Globicephala melas*.

### Outline extraction

Photographs of the fin platform and cross-sections were imported in AutoCAD software and calibrated using ruler marks. The fin planform outline and cross-sections were drawn manually using cubic B-splines tool in AutoCAD. As the fin cross-sections usually perform certain bend, the linear approximation procedure was applied to straighten the outline. The outline was divided into two parts using extreme points of the leading and trailing edges. 100 points were placed on each part in equal interval. Each couple of opposite points was joined by the complementary segment. Then the middle line passing through the middle points of all complementary segments was drawn. Coordinates of the middle and end points of each segment as well as length of the middle line were used for calculation of straightened outline’s coordinates.

### Wing and airfoil parameters of the fins

Following wing parameters were measured and calculated on the images of fins (Fig 5):

1. Fin span *S*, cm, measured from the fin base to the fin tip.
2. Basal length *BL* of the fin, cm, measured as length of the line parallel to the long axis of the body and starting from point of maximal curvature on the leading edge.
3. Leading edge length *L*, cm, measured from point of maximal curvature on the leading edge to the fin tip.
4. Fin area *A*, cm^2^, calculated with projection of the fin on the plane.
5. Angle of sweep *Λ*, degree, measured as angle between a perpendicular to root chord at the bottom of the fin and one-quarter chord position (Fig 6).
6. Aspect ratio *AR*, calculated as *S*^2^/*A*.
7. Canting index *CI*, calculated by the formula [8]:

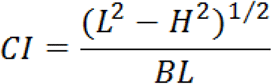

where *L* – leading edge length, *H* – span of the fin, and *BL* – basal length of the fin.

**Fig 5.**
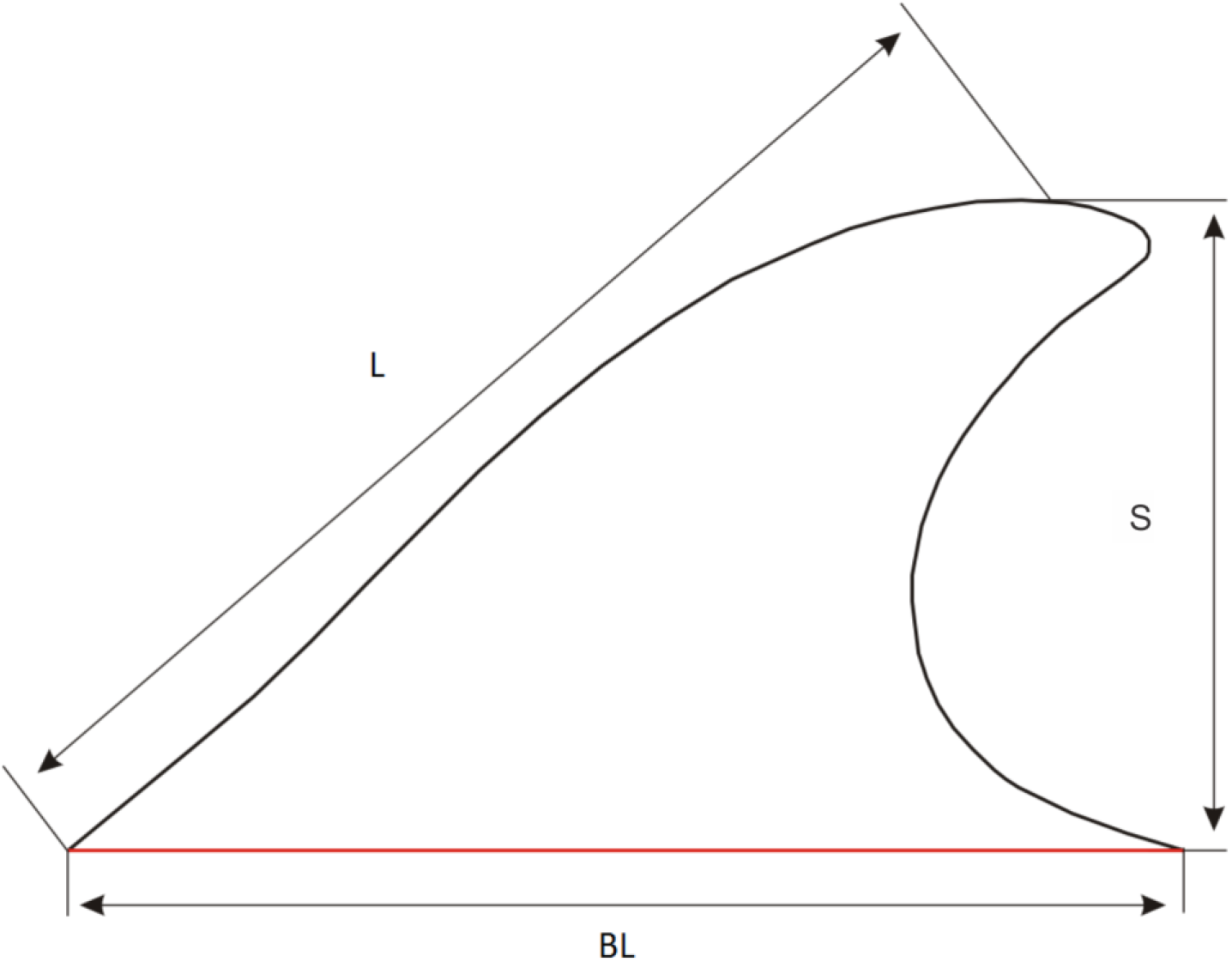
Basic measurements of fins.

**Fig 6.**
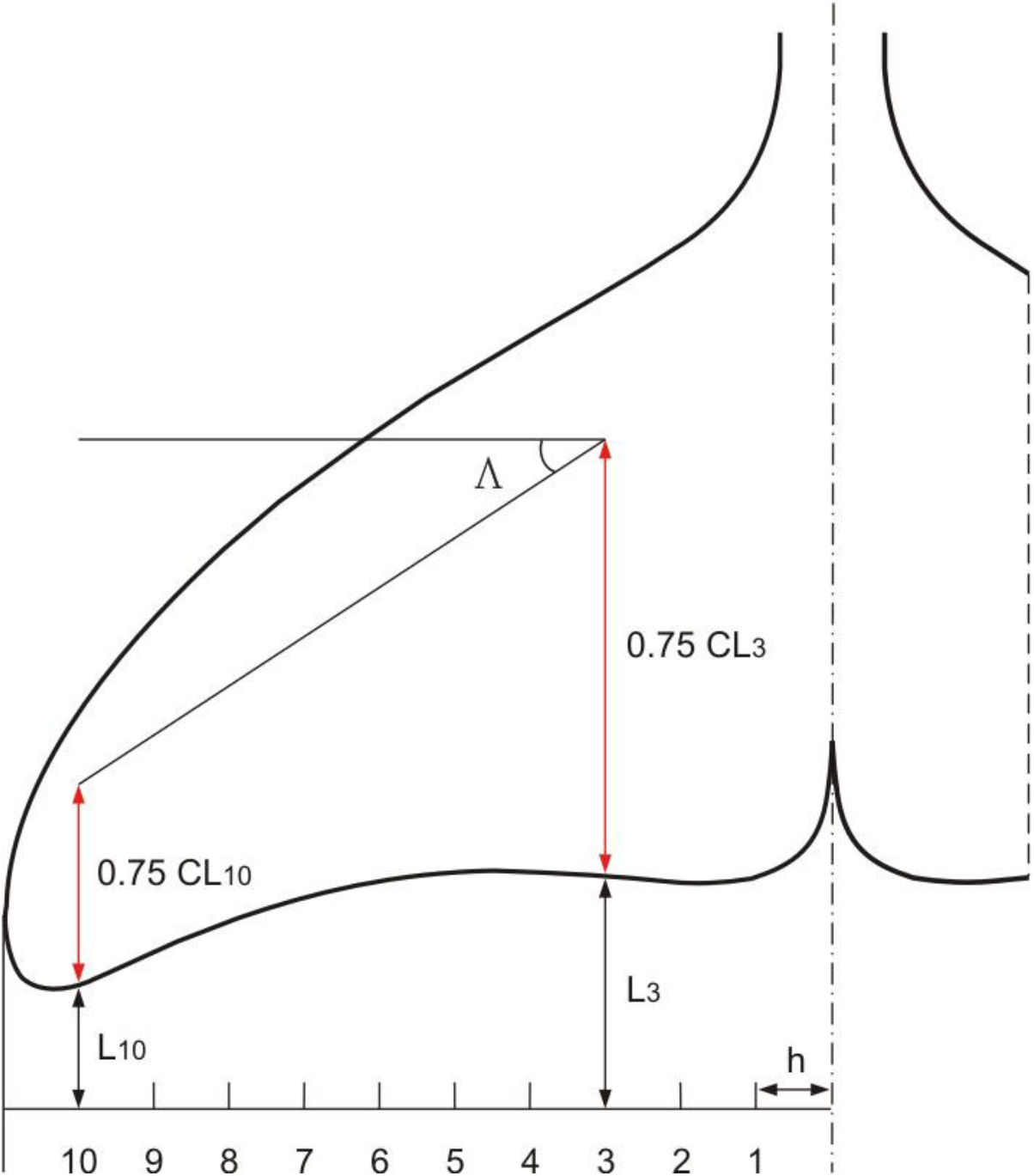
Scheme of measurement of sweep angle Λ on fins.

On straightened outlines of the cross-sections following airfoils parameters were measured and calculated (Fig 7):

1. Chord length *CL*, mm.
2. Maximal thickness *MT*, mm
3. Position of maximal thickness *PMT*, measured from the leading edge, mm.
4. Leading edge radius *r*, mm

**Fig 7.**
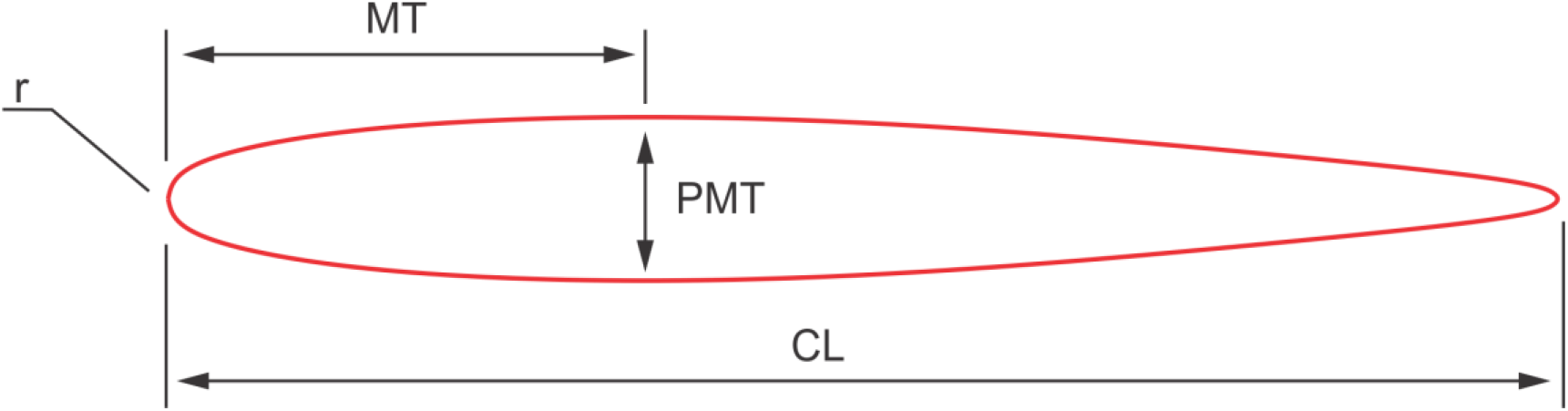
Scheme of the airfoil parameters measured on the cross-sections of fins. CL – chord length, MT – maximal thickness, PMT – position of maximal thickness, r – leading edge radius.

### Hydrodynamic performance of the fins cross-sections

Cross-sections of fins were analyzed with the DesignFOIL™ computer fluid dynamics software (DreeseCode Software). The water flow around the cross-sections and similar airfoils was simulated with a panel method. DesignFOIL™ software breaks the airfoil into many panels and forces the velocity at each panel to be tangential to that surface. Conglomerating all of these velocities leads to the velocity distribution and therefore the pressure coefficient distribution. The laminar flow portion of the boundary layer solver is based on the approximation von Karman and Pohlhauson method [17]. The turbulent flow is modeled on the approximation Buri method [45].

Reynolds number (*Re*), pressure distribution along the fin cross-sections, extent of the laminar, transition, and turbulent regions as well as lift coefficient *Cl*, drag coefficient *Cd*, moment coefficient *Cm*, and pressure coefficient *Cp* was calculated both for the cruising and burst speed of swimming assumed 2 m/sec and 8 m/sec respectively. All calculations were done for zero angle of attack α formed by the chord of a cross-section and the direction of the flow. Except of example below, all data in this study are referred to the animal gliding rectilinearly with constant speed. For the tail fluke cross-sections calculation of the hydrodynamic parameters was done out for the harbor porpoise, common dolphin, and bottlenose dolphin.

For comparison purpose, *Cd, Cl*, and *Cm* of the cross-sections taken at the bottom, mid-span, and the top of the tail flukes of one specimen of common dolphin and dorsal fin of one specimen of Atlantic white-beaked dolphin were calculated for the range of α from 0 to 20° and placed in drag polar diagram. Span-wise lift distribution on the cross-sections of the fluke of the common dolphin and dorsal fin of the Atlantic white-beaked dolphin was calculated as *CL*Cl_max_*, where *CL*, mm is a chord length of the symmetrical cross-sections, and *Cl_max_* is maximal lift coefficient calculated for the cross-section.

## Acknowledgements

Authors thank to the collaborators of LIENSS Institute of the University of La Rochelle, L’Observatoire PELAGIS UMS 3462, Museum of Natural History of the Faroe Islands, and ITAW Institute of the University of Hannover for their help and huge efforts in handling and morphometric measurements of cetaceans.

## Supporting information

**S1 Fig. Span-wise distribution of the chord length CL of the dorsal fin cross-sections, means ± SD.**

**S2 Fig. Span-wise distribution of the maximal thickness MT of the dorsal fin cross-sections, means ± SD.**

**S3 Fig. Span-wise distribution of the position of maximal thickness PMT of the dorsal fin cross-sections, means ± SD.**

**S4 Fig. Span-wise distribution of the relative maximal thickness MT, %CL of the dorsal fin cross-sections, means ± SD.**

**S5 Fig. Span-wise distribution of the relative position of maximal thickness PMT, %CL of the dorsal fin cross-sections, means ± SD.**

**S6 Fig. Span-wise distribution of the relative leading edge radius r, %CL of the dorsal fin cross-sections, means ± SD.**

**S7 Fig. Span-wise distribution of the chord length CL of the tail flukes cross-sections, means ± SD.**

**S8 Fig. Span-wise distribution of the maximal thickness MT of the tail flukes cross-sections, means ± SD.**

**S9 Fig. Span-wise distribution of the position of maximal thickness PMT of the tail fluke cross-sections, means ± SD.**

**S10 Fig. Span-wise distribution of the relative maximal thickness MT, %CL of the tail fluke cross-sections, means ± SD.**

**S11 Fig. Span-wise distribution of the relative position of maximal thickness PMT, %CL of the tail fluke cross-sections, means ± SD.**

**S12 Fig. Span-wise distribution of the relative leading edge radius r, %CL of the tail flukes cross-sections, means ± SD.**

**S13 Fig. Span-wise distribution of the drag coefficient of the dorsal fin cross-sections calculated at 2 m/sec, means ± SD.**

**S14 Fig. Span-wise distribution of the drag coefficient of the dorsal fin cross-sections calculated at 8 m/sec, means ± SD.**

**S15 Fig. Span-wise distribution of the relative laminar region LR, %CL of the dorsal fin cross-sections calculated at 2 m/sec, means ± SD.**

**S16 Fig. Span-wise distribution of the relative laminar region LR, %CL of the dorsal fin cross-sections calculated at 8 m/sec, means ± SD.**

**S17 Fig. Span-wise distribution of the drag coefficient of the tail flukes cross-sections calculated at 2 m/sec, means ± SD.**

**S18 Fig. Span-wise distribution of the drag coefficient of the tail flukes cross-sections calculated at 8 m/sec, means ± SD.**

**S19 Fig. Span-wise distribution of the relative laminar region LR, %CL of the tail fluke cross-sections calculated at 2 m/sec, means ± SD.**

**S20 Fig. Span-wise distribution of the relative laminar region LR, %CL of the tail fluke cross-sections calculated at 8 m/sec, means ± SD.**

**S21 Fig. Drag polar diagram of lift CL vs drag Cd I.** Calculated for the cross-sections taken at the bottom (dark blue), mid-span (light blue) and top (black) of the dorsal fin of Atlantic white-beaked dolphin at simulated swimming speed 2 m/sec.

**S22 Fig. Drag polar diagram of lift CL vs drag Cd II.** Calculated for the cross-sections taken at the bottom (dark blue), mid-span (light blue) and top (black) of the dorsal fin of Atlantic white-beaked dolphin at simulated swimming speed 8 m/sec.

**S23 Fig. Drag polar diagram of lift CL vs drag Cd III.** Calculated for the cross-sections taken at the bottom (dark blue), mid-span (light blue) and top (black) of the tail fluke of common dolphin at simulated swimming speed 2 m/sec.

**S24 Fig. Drag polar diagram of lift CL vs drag Cd VI.** Calculated for the cross-sections taken at the bottom (dark blue), mid-span (light blue) and top (black) of the tail fluke of common dolphin at simulated swimming speed 8 m/sec.

**S25 Fig. Span-wise lift distribution.** Calculated for the cross-sections of the fluke of the common dolphin (blue) and dorsal fin of the Atlantic white-beaked dolphin (green) at simulated swimming speed 2 and 8 m/sec.

## Notes

### Competing Interest Statement

The authors have declared no competing interest.

