## Supplemental Figures S1-S25 for "Form, function, and divergence of a generic fin shape in small cetaceans"

### Supporting information

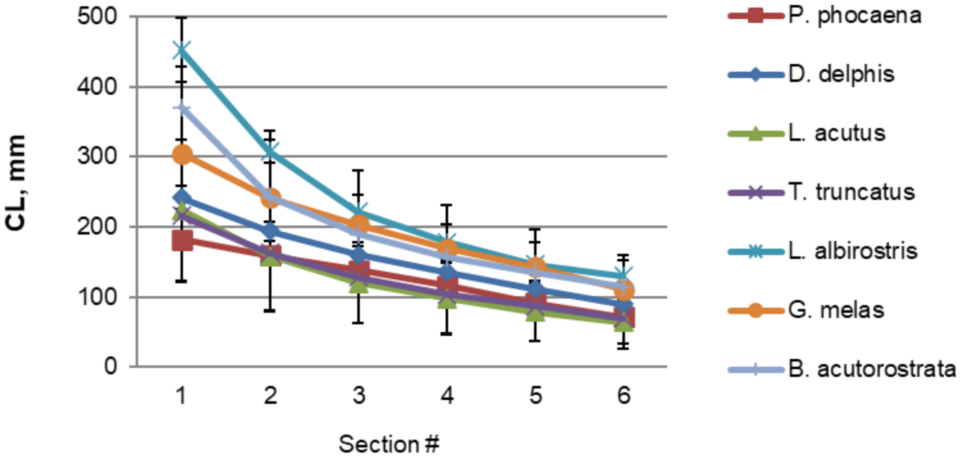


**S1 Fig.**

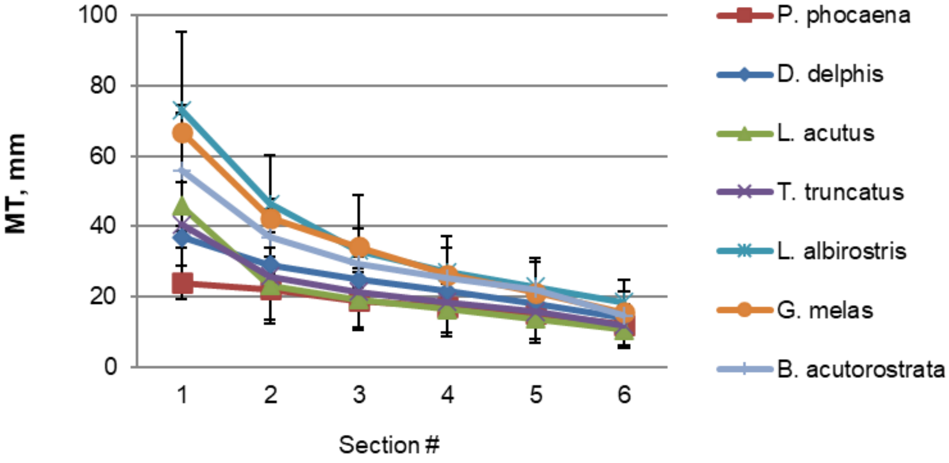


**S2 Fig.**

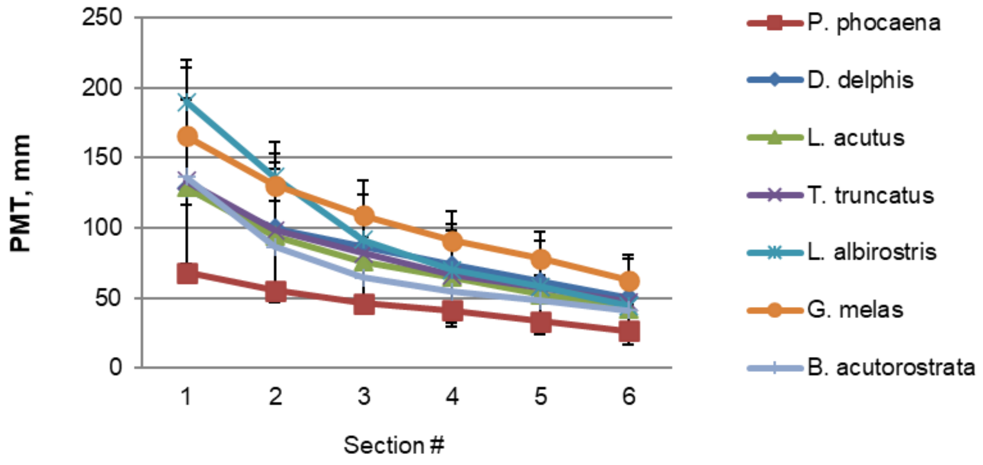

PMT, mm
250
200
150
100
50
0
1
2
3
4
5
6
Section #
P. phocaena
D. delphis
L. acutus
T. truncatus
L. albirostris
G. melas
B. acutorostrata


**S3 Fig.**

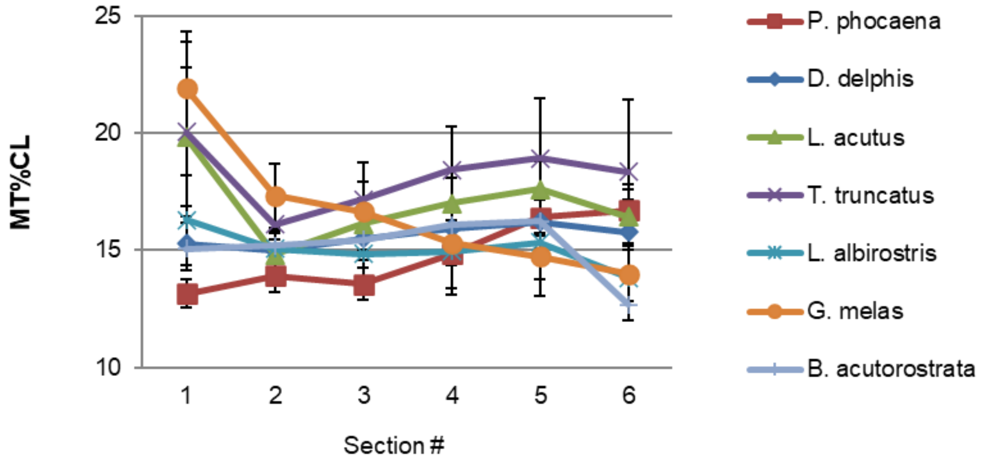

MT%CL
25
20
15
10
1
2
3
4
5
6
Section #
P. phocaena
D. delphis
L. acutus
T. truncatus
L. albirostris
G. melas
B. acutorostrata


**S4 Fig.**

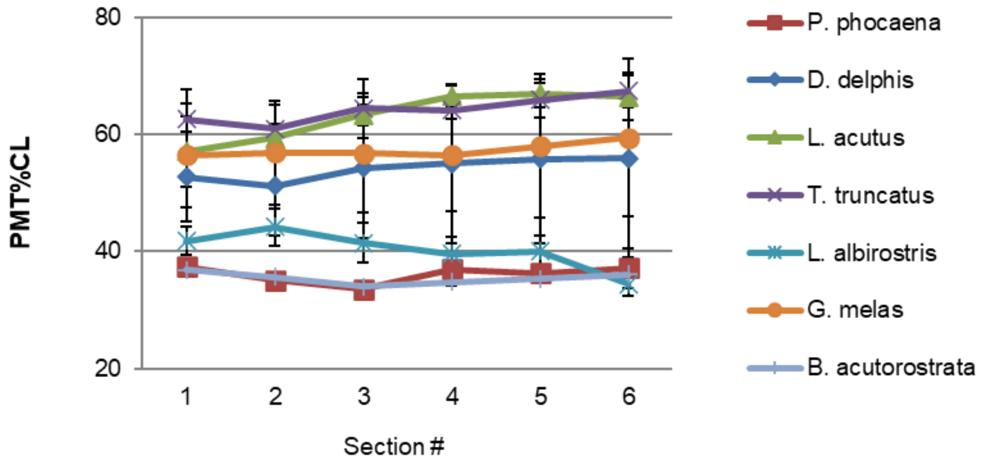
PMT%CL
80
60
40
20
1
2
3
4
5
6
Section #
P. phocaena
D. delphis
L. acutus
T. truncatus
L. albirostris
G. melas
B. acutorostrata

**S5 Fig.**

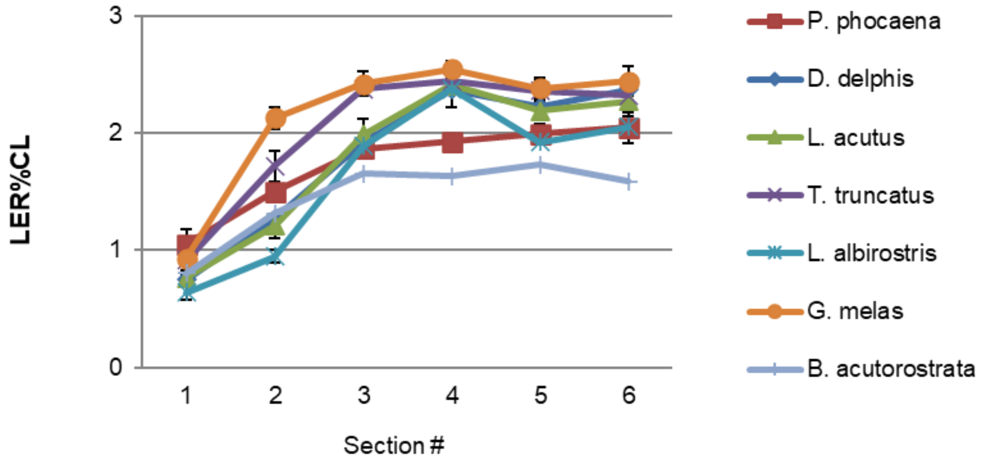
LER%CL
3
2
1
0
1
2
3
4
5
6
Section #
P. phocaena
D. delphis
L. acutus
T. truncatus
L. albirostris
G. melas
B. acutorostrata

**S6 Fig.**

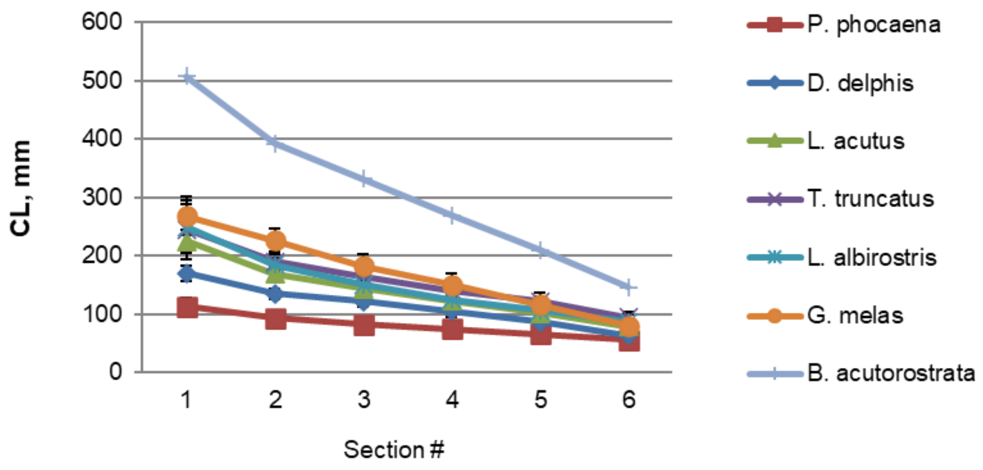
600
500
400
300
200
100
0
CL, mm
1
2
3
4
5
6
Section #
P. phocaena
D. delphis
L. acutus
T. truncatus
L. albirostris
G. melas
B. acutorostrata

**S7 Fig.**

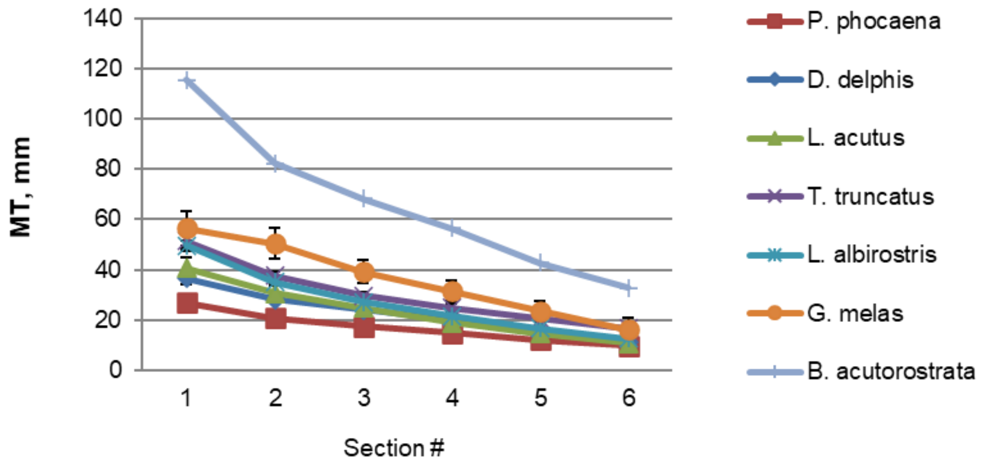
140
120
100
80
60
40
20
0
MT, mm
1
2
3
4
5
6
Section #
P. phocaena
D. delphis
L. acutus
T. truncatus
L. albirostris
G. melas
B. acutorostrata

**S8 Fig.**

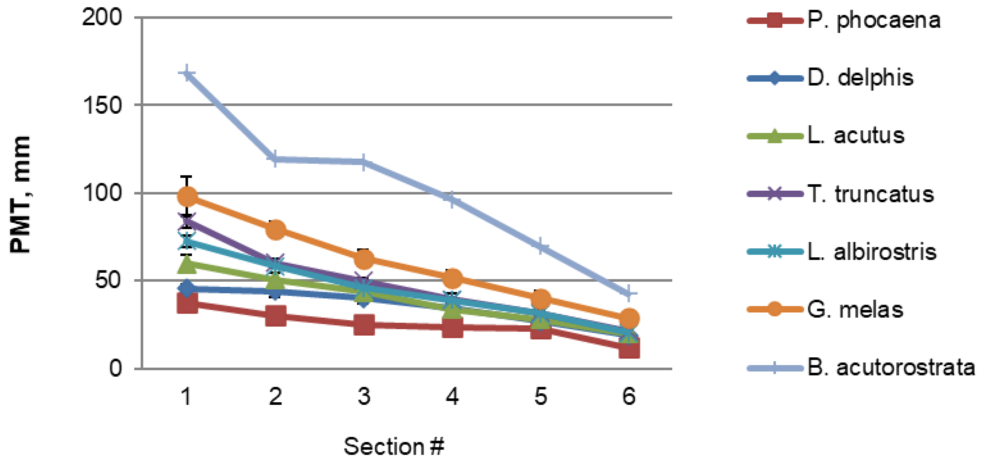


**S9 Fig.**

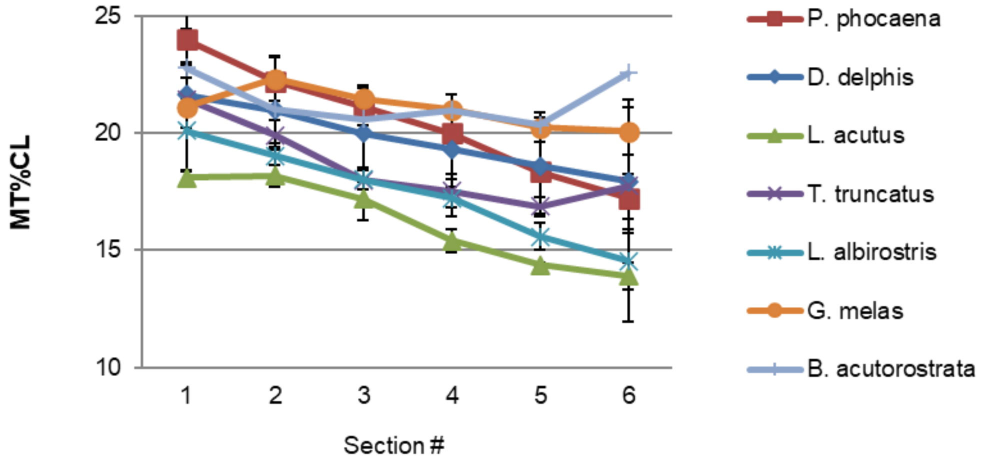


**S10 Fig.**

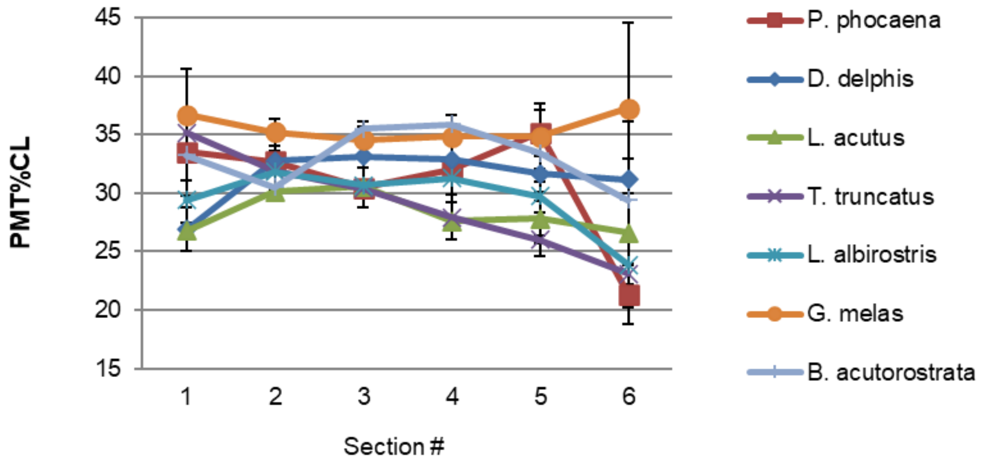


**S11 Fig.**

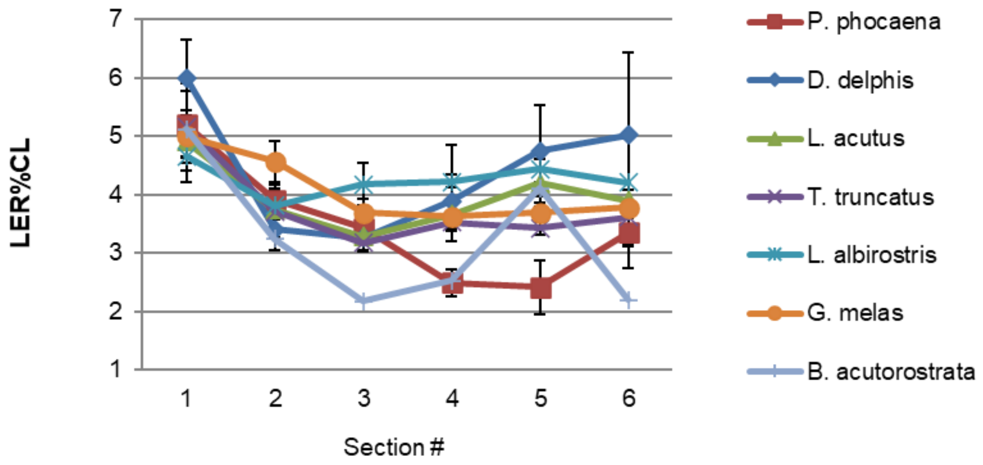


**S12 Fig.**

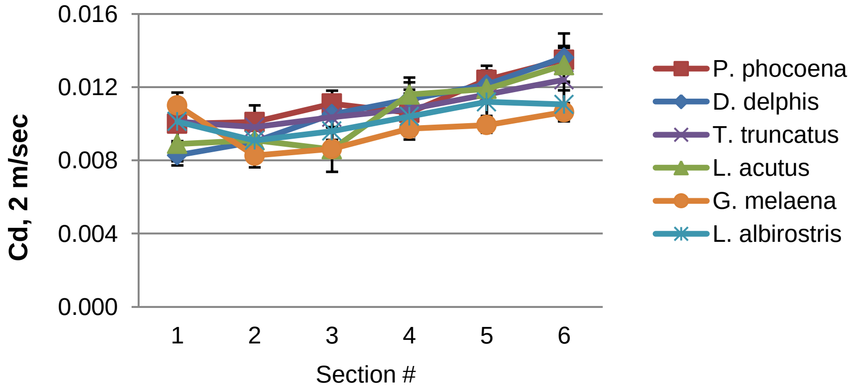


**S13 Fig.**

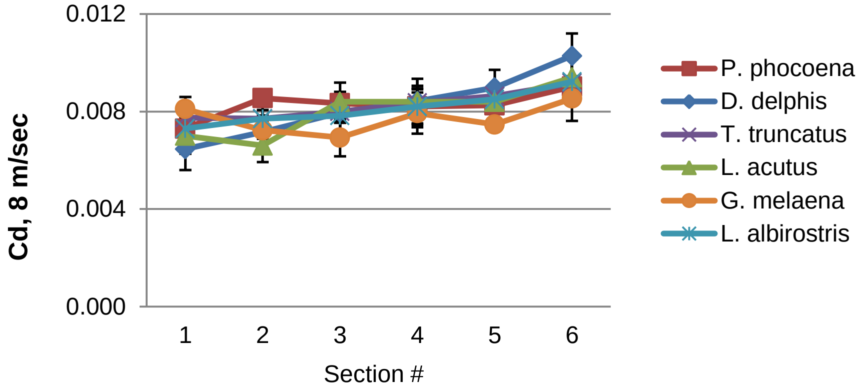


**S14 Fig.**

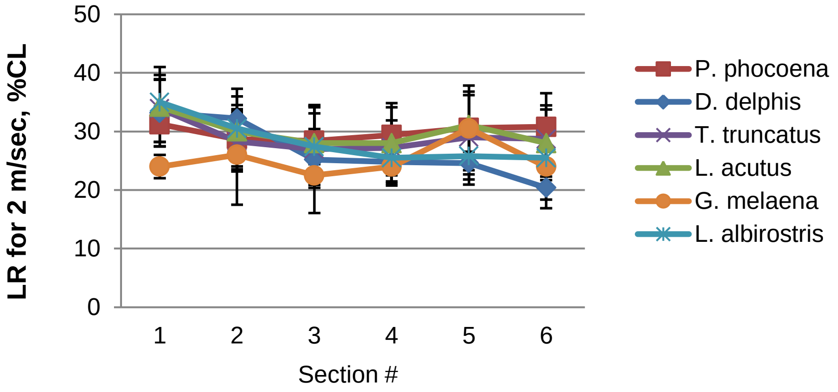


**S15 Fig.**

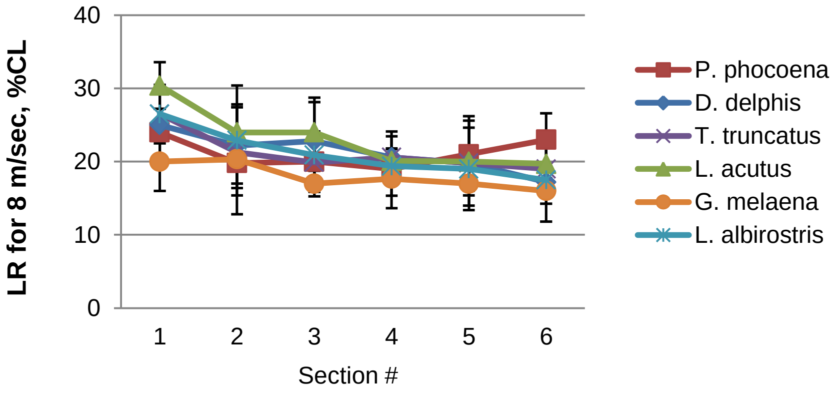


**S16 Fig.**

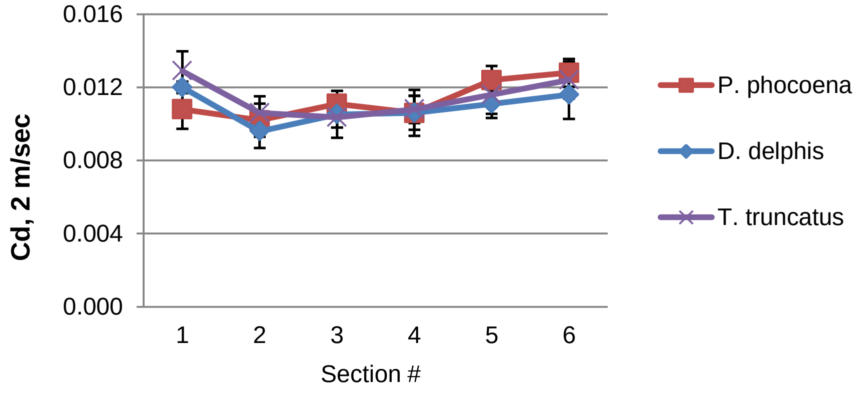


**S17 Fig.**

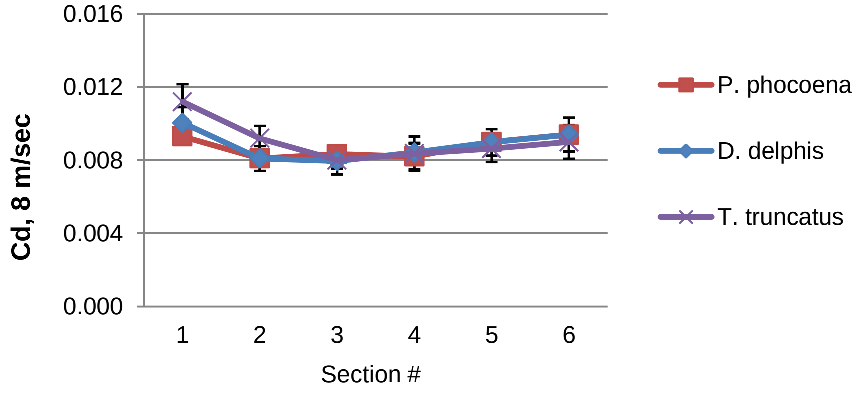


**S18 Fig.**

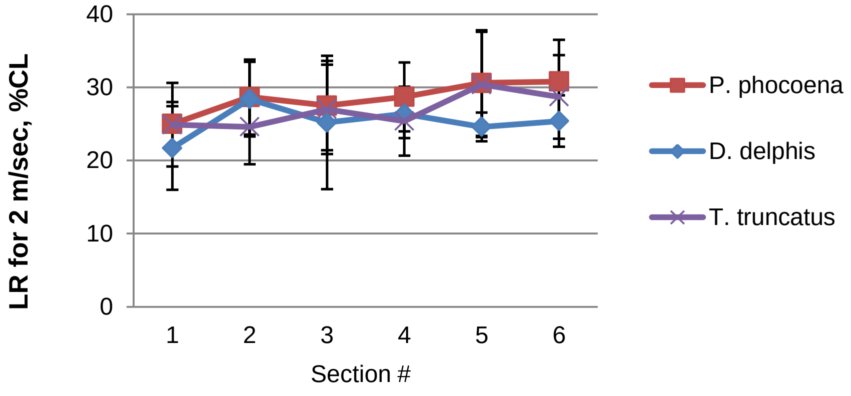


**S19 Fig.**

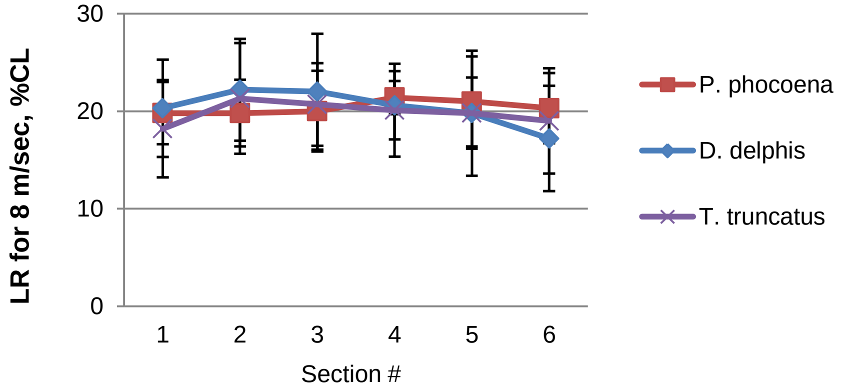


**S20 Fig.**

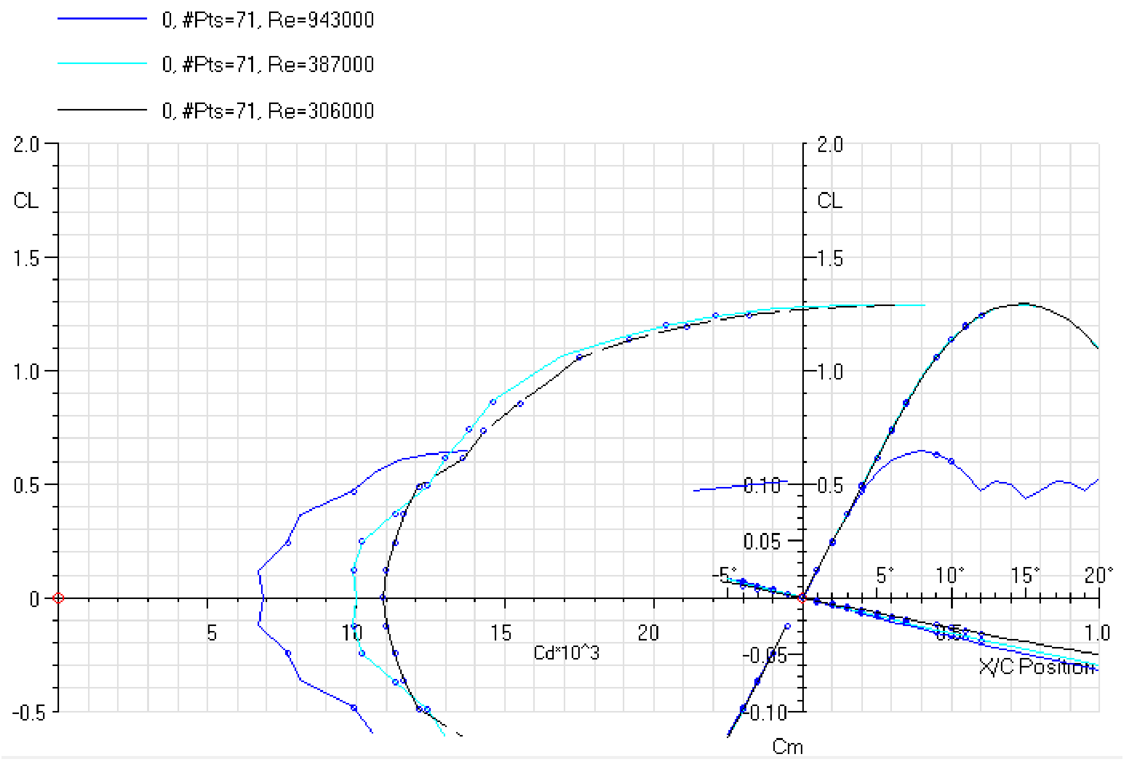
0, #Pts=71, Re=943000
0, #Pts=71, Re=387000
0, #Pts=71, Re=306000
CL
Cd*10^3
Cm
X/C Position

**S21 Fig.**

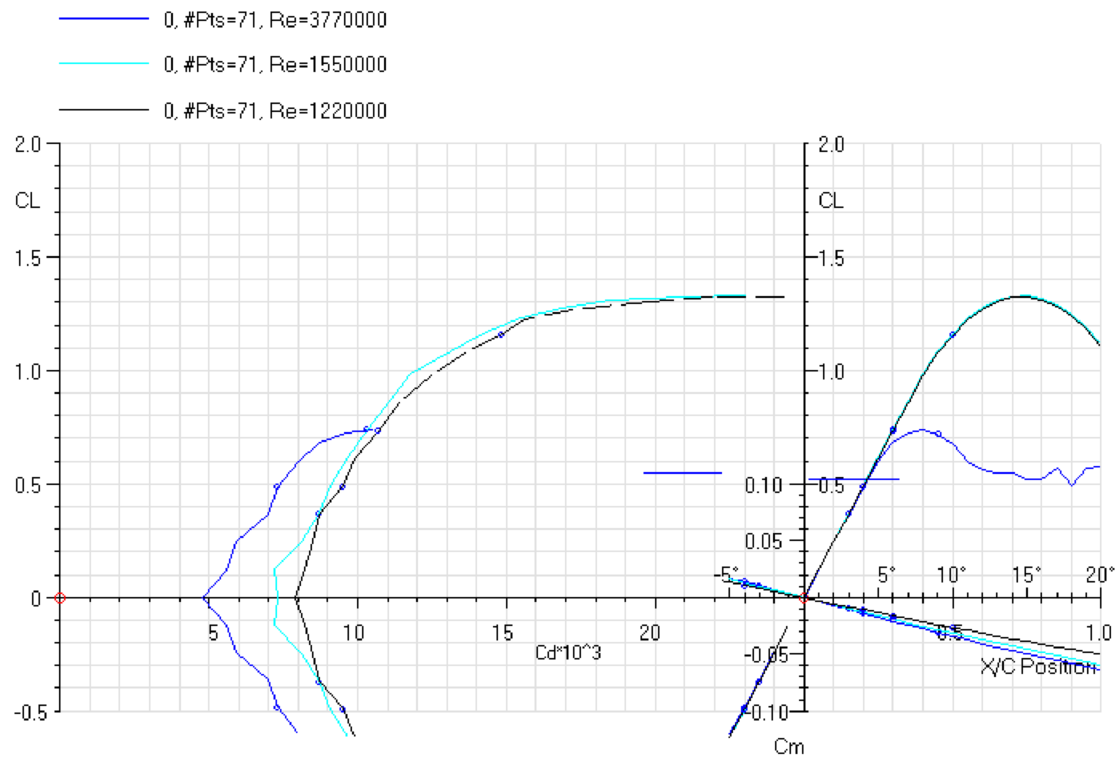
0, #Pts=71, Re=3770000
0, #Pts=71, Re=1550000
0, #Pts=71, Re=1220000
CL
Cd*10^3
Cm
X/C Position

**S22 Fig.**

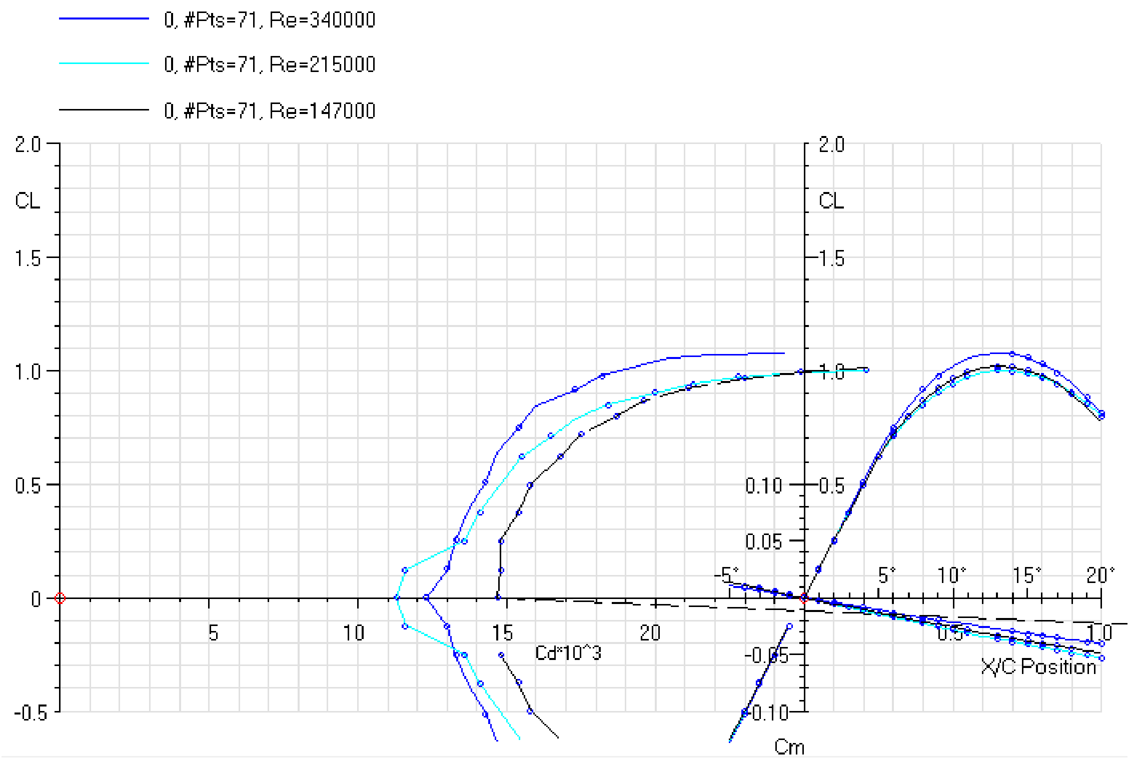
0, #Pts=71, Re=340000
0, #Pts=71, Re=215000
0, #Pts=71, Re=147000
CL
Cd*10^3
Cm
X/C Position
2.0
1.5
1.0
0.5
0
-0.5
5
10
15
20
0.10
0.05
-0.05
-0.10
-5°
5°
10°
15°
20°
0.5
1.0

**S23 Fig.**

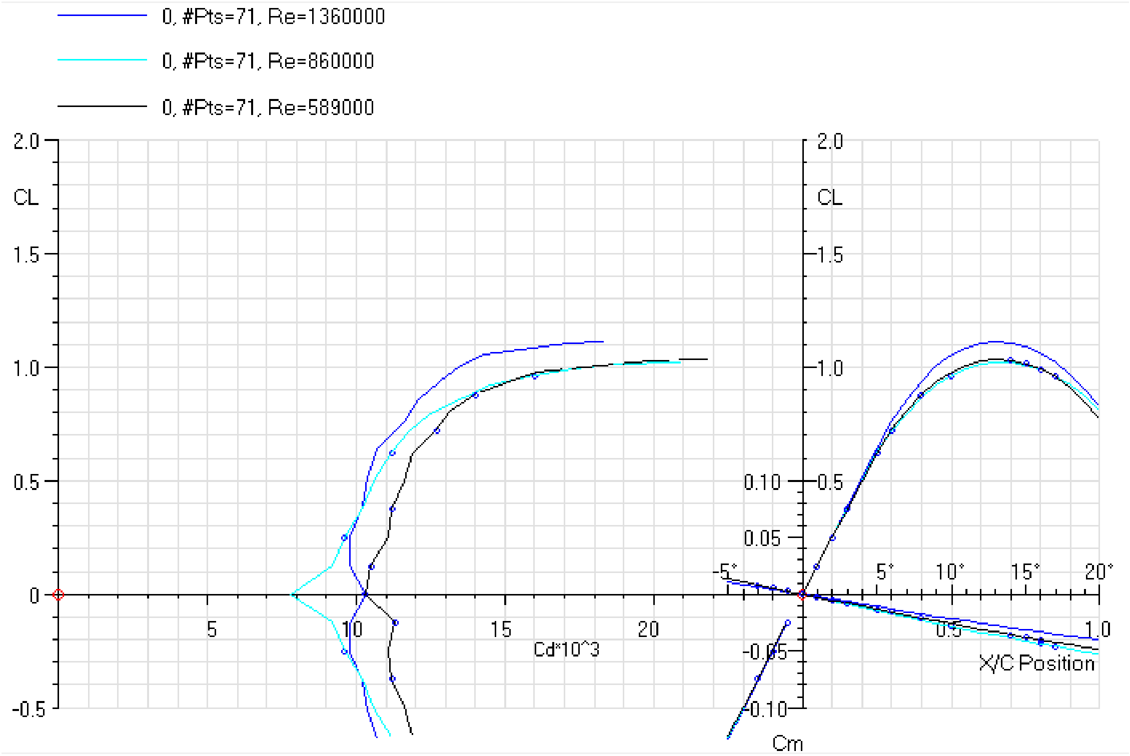
0, #Pts=71, Re=1360000
0, #Pts=71, Re=860000
0, #Pts=71, Re=589000
CL
2.0
1.5
1.0
0.5
0
-0.5
5
10
15
20
Cd*10^3
0.10
0.05
-0.05
-0.10
Cm
-5°
5°
10°
15°
20°
0.5
1.0
X/C Position

**S24 Fig.**

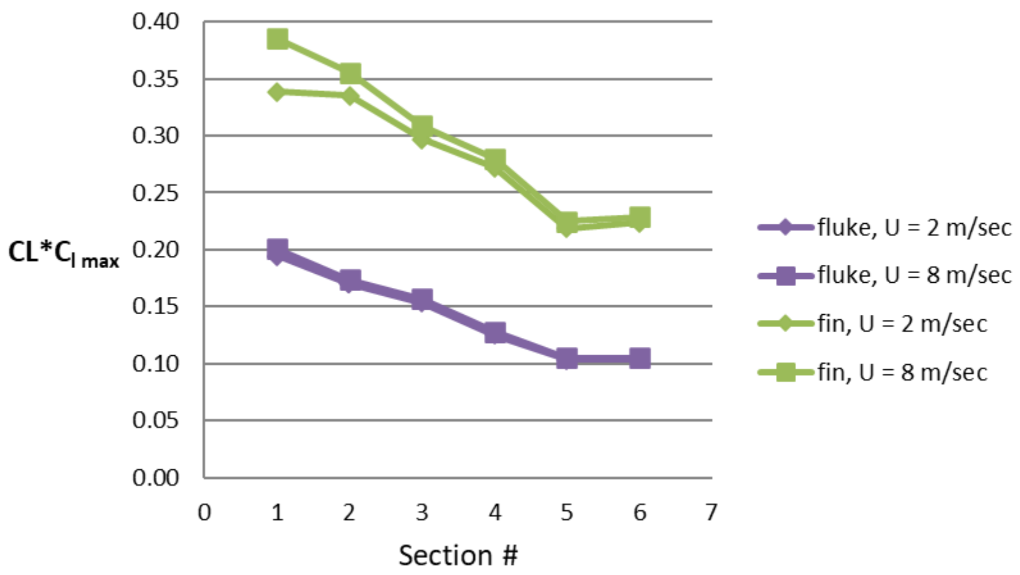
CL*Cl max
0.40
0.35
0.30
0.25
0.20
0.15
0.10
0.05
0.00
0
1
2
3
4
5
6
7
Section #
fluke, U = 2 m/sec
fluke, U = 8 m/sec
fin, U = 2 m/sec
fin, U = 8 m/sec

**S25 Fig.**
